# Multiomics Approach to Regionally Profile Zinc-Driven Host-Gut Microbiome Interactions in the Intestinal Tract

**DOI:** 10.1101/2025.07.02.662895

**Authors:** S Sastry, SB Mitchell, A Glick, TL Thorn, TB Aydemir

**Affiliations:** Cornell University, Division of Nutritional Sciences, Ithaca, NY 14853

**Keywords:** Zinc Deficiency, Spatial Regulation, Transcriptome, Inflammatory Diseases, Metabolic Diseases

## Abstract

Zinc deficiency (ZnD) is a major risk factor for metabolic and inflammatory diseases associated with gut microbial alterations, such as obesity, type 2 diabetes, and inflammatory bowel disease. Despite its importance, there are no established dietary recommendations for zinc supplementation in individuals with these conditions, except in cases of severe diarrhea in children. This gap stems from inconsistent outcomes in zinc supplementation trials, suggesting an incomplete understanding of zinc-mediated spatial and temporal regulation within the host-microbiome interface. This study employed a multiomics approach to investigate zinc-driven host-gut microbial interactions in the intestinal tract of mice fed zinc-adequate or ZnD diets. Radio tracing and metallomics analyses uncovered differential zinc abundance across intestinal tissues, with conventionalized germ-free mice displaying significantly lower zinc levels than germ-free mice, highlighting the reciprocal regulation of zinc between host tissues and gut microbes. Transcriptomic analyses revealed region-specific effects of ZnD, including altered energy metabolism and apoptosis in the small intestine, and impaired barrier function and redox processes in the colon. Metagenomics profiling revealed that ZnD reduced microbial diversity in the small intestine, while the cecum and colon were protected from diversity loss but exhibited an increased abundance of pathogenic bacteria. Correlation analyses linked tissue-specific host gene expression to shifts in microbial populations, identifying potential microbial mediators of host transcription changes under ZnD. Collectively, these findings emphasize the critical role of zinc in spatially regulating host-microbiome interactions, advancing our understanding of region-specific impacts of ZnD on the GI tract and disease risk.

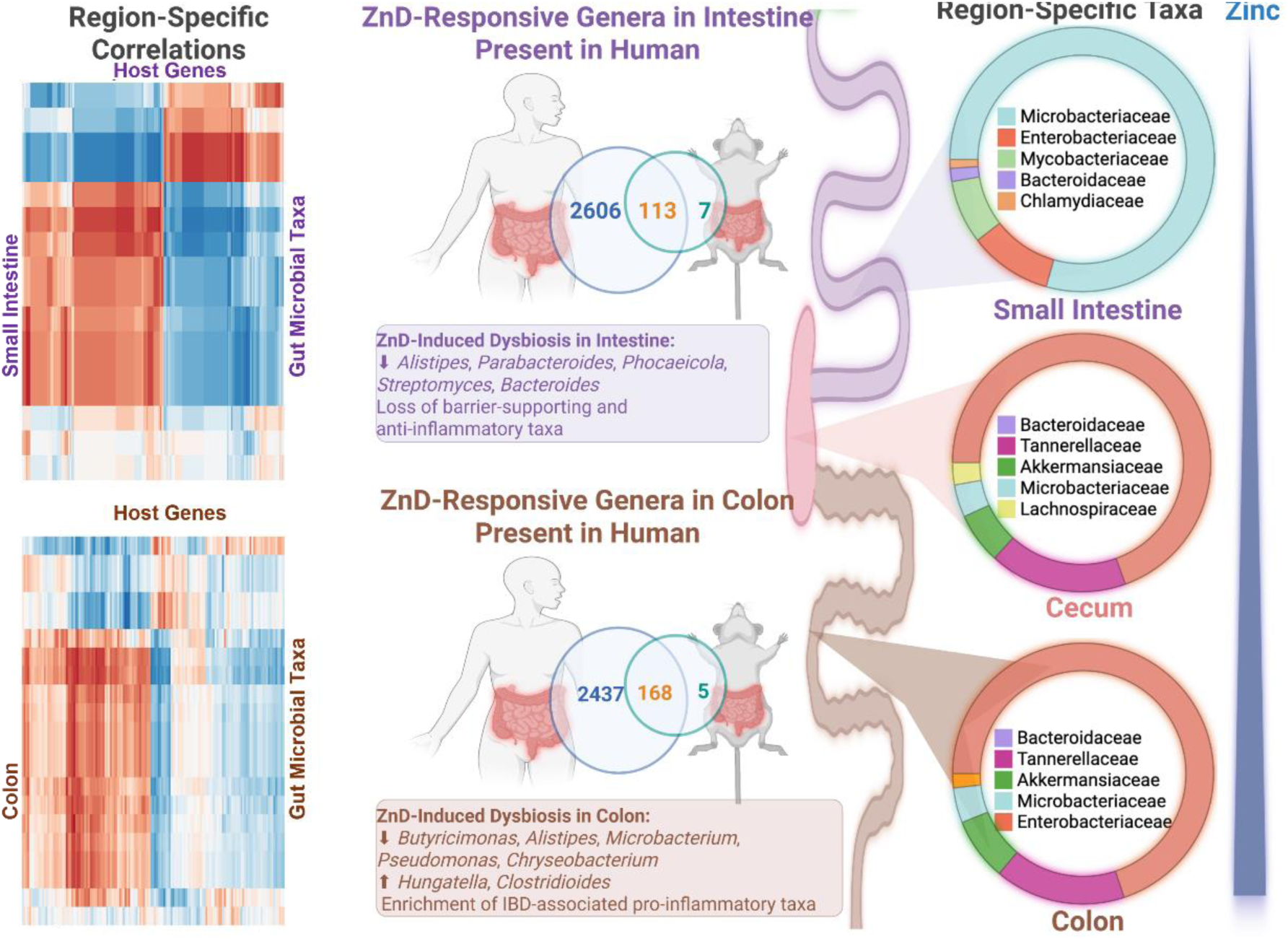

## Introduction

Zinc is an essential micronutrient required by virtually all living organisms, including humans and microbes, and plays a pivotal role in host-microbiome interactions along the gastrointestinal (GI) tract. ^1–4^ This coordination is critical for maintaining both metabolic and inflammatory homeostasis. Zinc deficiency (ZnD) has been identified as a significant risk factor for a range of human diseases associated with gut microbial dysbiosis, including obesity, type 2 diabetes mellitus, and inflammatory bowel disease.^5–7^ Its importance is particularly well-documented in cases of severe childhood diarrhea, where zinc supplementation is routinely recommended as a therapeutic intervention. ^8^ However, beyond this context, universally established dietary reference intakes and guidelines for zinc supplementation in individuals with metabolic or inflammatory conditions associated with ZnD remain unavailable. The absence of standardized recommendations is largely driven by inconsistent outcomes from zinc supplementation trials, ^9–19^ highlighting the complex and context-specific roles of zinc in regulating host-microbiome dynamics. These inconsistencies point to a critical knowledge gap in understanding how zinc mediates both spatial and temporal interactions at the host-gut microbiome interface.

Zinc homeostasis is tightly regulated in the gastrointestinal tract through dietary zinc absorption, reabsorption of endogenous zinc secretions, and zinc excretion.^20^ Dietary zinc absorption occurs along the entire intestinal tract, with the proximal intestine serving as the dominant site of uptake. ^21,22^ Endogenous zinc is lost via pancreatic secretions, sloughing of intestinal epithelial cells, and transepithelial flux of zinc from enterocytes into the lumen.^23,24^ Although most endogenous zinc excreted into the GI tract is lost in feces, a portion is reabsorbed to sustain systemic zinc balance. This dynamic regulation results in region-specific variations in zinc abundance within distinct gastrointestinal tissues and luminal contents, with potential implications for the spatial organization and composition of gut microbiota; however, the impact of these spatial variations in zinc metabolism on gut microbial communities remains poorly understood.

The gastrointestinal tract itself is a spatially dynamic system, with region-specific nutrient gradients, oxygenlevels, microbial communities, and immune responses ^25^ that collectively define intestinal health and disease susceptibility. Emerging evidence highlights the compositional and functional distinctions between the microbiota in the intestine and colon, which are shaped by localized host factors and environmental cues.^26–29^ These spatial differences are particularly relevant in the context of metabolic and inflammatory conditions, where ZnD and gut dysbiosis frequently coexist. Despite this interplay, the mechanisms through which zinc shapes spatial host-microbiome dynamics across the gastrointestinal tract remain elusive.

To address this gap, the present study employs integrative multiomics approaches to regionally profile gastrointestinal zinc distribution, host transcriptomic changes, gut microbiome composition, and their interrelationships in mice maintained on zinc-adequate or zinc-deficient diets. By unraveling how regional differences in zinc availability mediate host-microbiome interactions, this study aims to provide novel insights into the mechanisms underlying dysbiosis-driven metabolic and inflammatory diseases, with the ultimate goal of informing therapeutic strategies targeting ZnD-associated conditions.

## Results

**Interplay between spatial gastrointestinal zinc distribution and gut microbiota.**

In this study, three strains of mice were used to examine the regional distribution of zinc in the gastrointestinal tract. Using radiotracer ^65^Zn, we measured radioactivity in gastrointestinal tissues, luminal content, peripheral tissues, and serum to analyze zinc transport dynamics (**Figure 1A**). Results revealed that when delivered orally, the colon displayed significantly lower levels of ^65^Zn compared to the proximal and distal segments of the small intestine, between which no significant difference was observed (**Figure 1B**). When delivered via subcutaneous injection, a gradual and significant decline in the amount of ^65^Zn from the proximal to distal intestine and finally to the colon was observed (**Figure 1B**). Consistent with these findings, the lumen of the colon retained significantly more ^65^Zn when the tracer was given orally and significantly less ^65^Zn when administered via injection compared to the lumen of the small intestine (**Figure 1C**). Regardless of route administration, we observed a significant correlation between the luminal zinc contents of the small intestine, suggesting tight regulation and equilibrium of zinc transport in both directions (**Supp. Figure 1A**). However, no such correlation was observed in the colon (**Supp. Figure 1B**).

**Figure 1.**
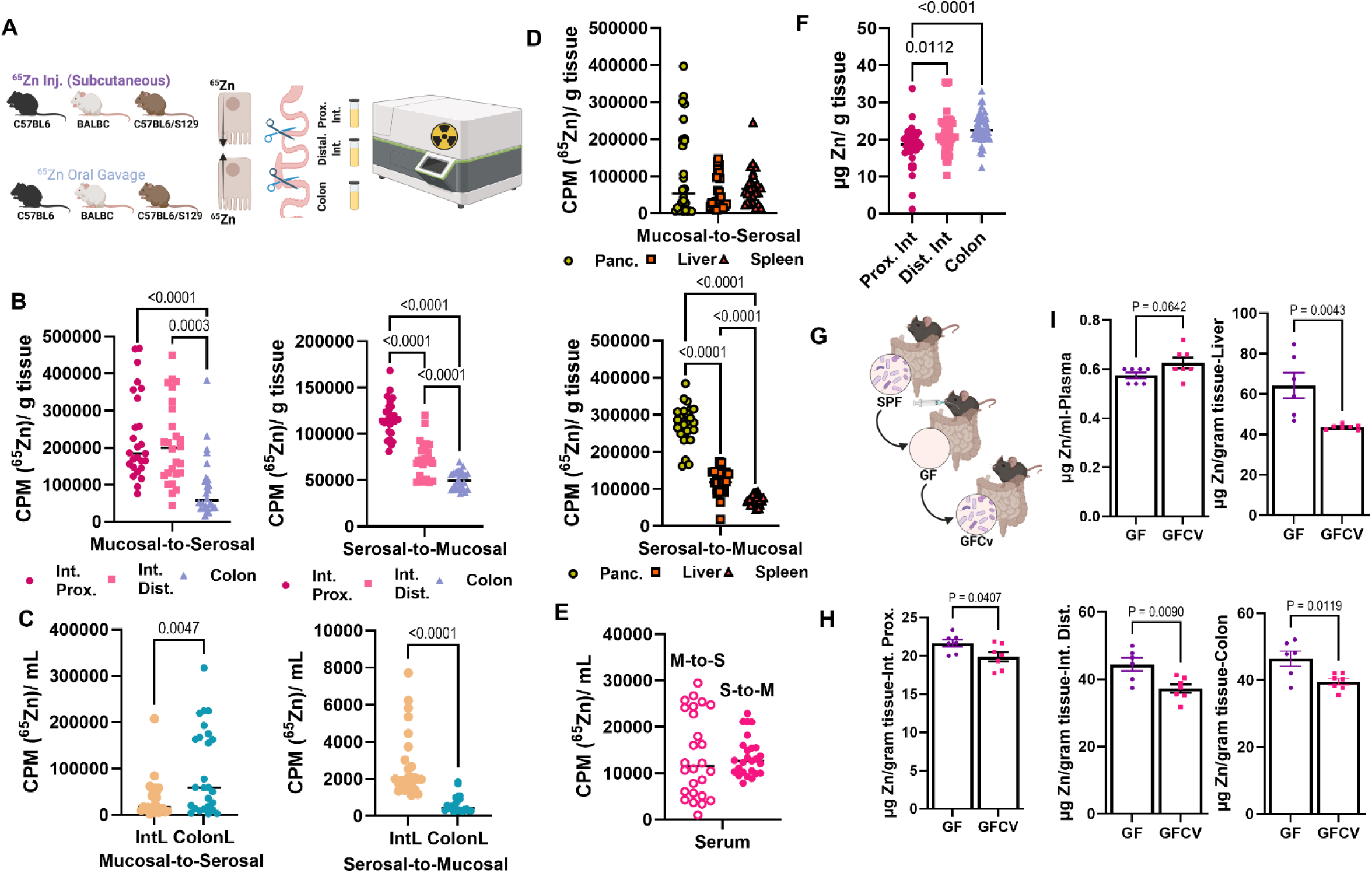
Gastrointestinal zinc distribution changes spatially and in response to microbiota. (A) Overview of study design. ^65^Zn radioisotope tracing was conducted across three strains of mice (C57BL6, BALBC, and C57BL6/S129) in both the serosal-to-mucosal (Subcutaneous injection) and mucosal-to-mucosal (Oral gavage) directions. Radioactivity was measured by gamma counting 3 hours after ^65^Zn administration in the proximal small intestine, distal small intestine, and colon to measure Zn distribution. (B-E) Counts per minute (CPMs) in gastrointestinal tissues (B), intestine and colon luminal content (C), peripheral tissues (Pancreas, Liver, Spleen) (D) and serum (E) following ^65^Zn administration. (F) Zn concentrations in gastrointestinal tissues as measured by microwave plasma atomic emission spectroscopy (MP-AES). (G) Feces from specific pathogen-free mice (SPF) was transplanted into germ-free mice (GF) by oral gavage. Resulting germ-free-conventionalized (GFCv) were compared to GF mice (n = 6-7). (H,I) Zn concentrations as measured by MP-AES in gastrointestinal tissues (H), plasma, and liver tissue (I) in GF and GFCv mice.

To evaluate the systemic impact of zinc transport dynamics, ^65^Zn levels were measured in peripheral tissues, including the pancreas, liver, and spleen (**Figure 1D**). The pancreas and liver, both known for their strong responsiveness to zinc, were included in the analysis, while the spleen served as a negative control. When ^65^Zn was administered orally, no differences in ^65^Zn accumulation were observed among the examined tissues. However, when ^65^Zn was delivered via subcutaneous injection, the highest levels of ^65^Zn were found in the pancreas, followed by the liver and then the spleen. These findings suggest that both the pancreas and liver play significant roles in maintaining systemic zinc homeostasis, working in coordination with the gastrointestinal tract. Interestingly, the spleen maintained the same level of ^65^Zn regardless of the administration route, supporting its role as a negative control in the study. Additionally, serum ^65^Zn levels remained unchanged irrespective of whether the tracer was administered orally or by injection (**Figure 1E**). Lastly, we measured the zinc concentrations in the sections of mouse intestinal tissues. We found significantly greater zinc concentrations in the colon than in the proximal intestine, with no differences between the proximal and distal intestine (**Figure 1F**).

We then tested the impact of gut microbiota on gastrointestinal and systemic zinc metabolism by using germ-free and conventionalized mice (**Figure 1G**). Significantly lower zinc levels were observed in both the small intestine and colon of conventionalized mice compared to germ-free mice (**Figure 1H**), suggesting the presence of zinc-acquiring gut microbes in both the intestine and colon. We also found significantly lower levels of hepatic zinc in the conventionalized mice, and a trend towards greater zinc concentrations in the plasma.

### The gastrointestinal tract harbors a taxonomically distinct microbial environment, with the most pronounced differences observed in bacterial communities

To elaborate upon the reciprocal relationship between zinc and microbes across intestinal regions, we analyzed microbial communities across the small intestine, cecum, and colon. Principal Coordinates Analysis (PCoA) based on Bray-Curtis dissimilarity revealed distinct clustering of microbial communities by intestinal region across all domains profiled using the Kraken2 PlusPF database, which includes bacterial, archaeal, viral, and fungal genomes.^30^ The small intestine consistently clustered separately from the cecum and colon, indicating a unique microbial composition in this region (**Figure 2A**). Given the dominance of bacteria, comprising a mean relative abundance of nearly 99% across all samples (**Figure 2B),** we further assessed distinct clustering of microbial communities by intestinal region based solely on bacterial reads. This separation was also pronounced in the bacterial community, where intestinal samples were clearly resolved from cecal and colonic communities along the primary coordinate (PC1, 86.1%) (**Figure 2C**). Fungal and viral community structures also showed some separation by location, but the effect was less prominent than in bacterial reads (**Supp. Figure S2A**), likely reflecting lower abundance or diversity in non-bacterial taxa.

**Figure 2.**
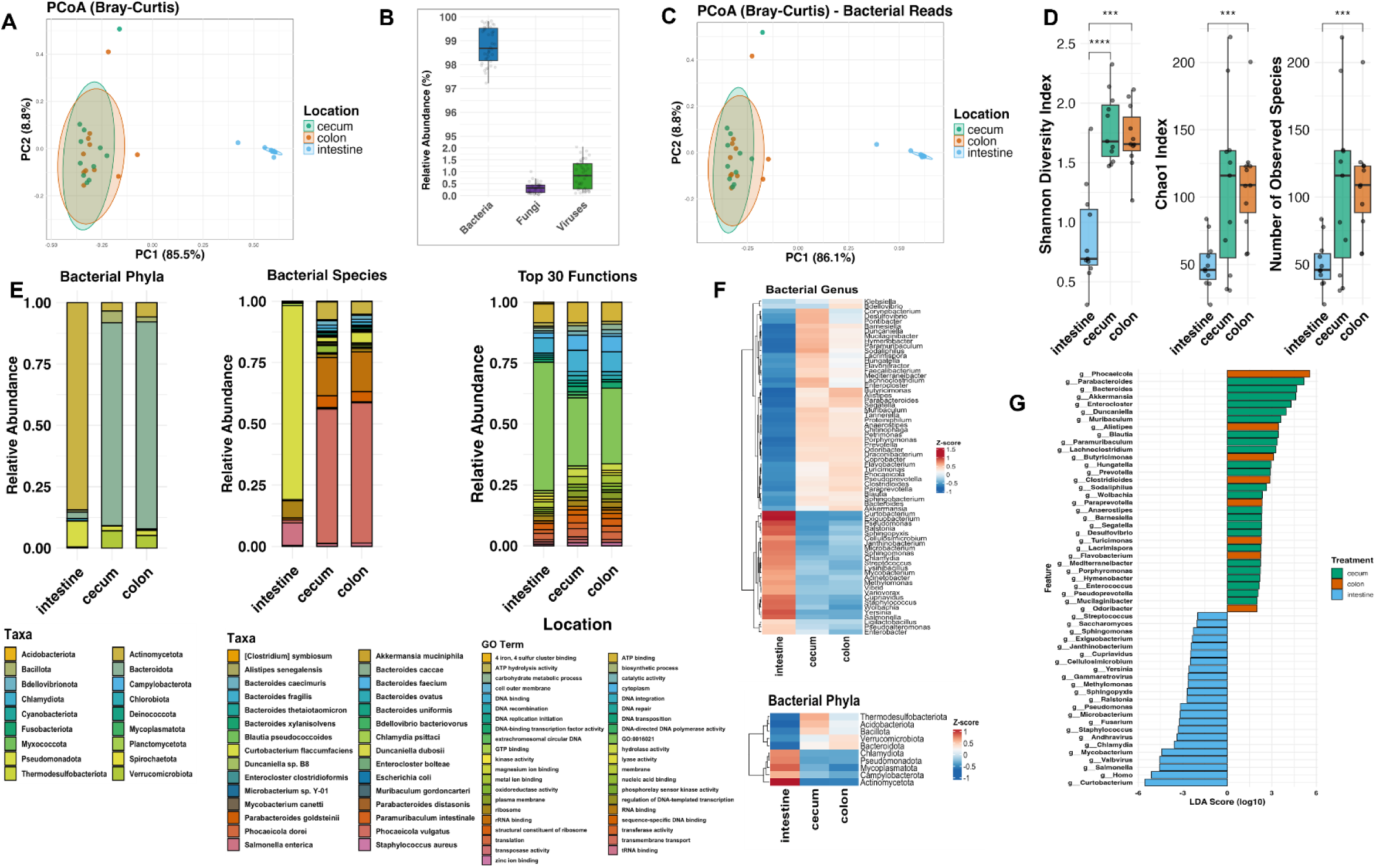
Region-specific shifts in gut microbial composition and function across the intestine, cecum, and colon. (A) Multikingdom community structure. PCoA of Bray–Curtis dissimilarities calculated from all classified reads (bacteria, fungi, viruses, archaea; n = 33 samples). Ellipses depict the 95 % confidence region for each intestinal location. The first two axes explain 85.5 % (PC1) and 8.8 % (PC2) of the variance. (B) Kingdom-level read distribution. Box-and-jitter plots show the proportion of bacterial, fungal and viral reads per sample after Kraken2/Bracken classification (C) Bacterial community structure. PCoA of Bray–Curtis dissimilarities computed from bacterial reads only. Ellipses depict the 95 % confidence region for each intestinal location. PC1 = 86.1 %, PC2 = 8.8 %. (D) α-diversity of bacterial reads. Shannon diversity, Chao1 richness, and observed species were estimated in phyloseq in R version 4.4.2. Boxes are median ± IQR; points are individual mice (intestine n = 11, cecum n = 11, colon n = 11). Pairwise Wilcoxon rank-sum tests with Benjamini–Hochberg correction: P < 0.001 (***) indicates significance for pairwise comparison. (E) Compositional and functional profiles. Bacterial phyla & species: Stacked bars display the mean relative abundance of phyla and species level bacterial reads. The 30 most abundant species in each intestinal region. Abundances were TSS-normalised within samples, then averaged by region. Top 30 GO terms: HUMAnN3 (v 3.8) gene-family counts were collapsed to GO terms, converted to counts-per-million, and the 30 most abundant functions per region were plotted. (F) Differentially abundant taxa. Heatmaps show Z-score-scaled log10 relative abundances of bacterial genera and phyla that were significantly associated with location in MaAsLin2 with a linear model that included intestinal region as a fixed effect. Taxa with FDR-adjusted p < 0.05 were considered significant. Comparisons were made relative to the small intestine. Rows are hierarchically clustered (Euclidean distance, complete linkage). (G) LEfSe analysis. Linear discriminant analysis effect-size (LEfSe) identified region-enriched bacterial genera (CPM normalization, α = 0.05 Kruskal–Wallis, LDA ≥ 2.0). Bars represent signed log10 LDA scores.

### Alpha diversity metrics reveal opposing regional trends for bacterial vs. non-bacterial taxa

Alpha Diversity across the small intestine, cecum, and colon was examined using three metrics: Shannon diversity, observed species richness, and Chao1 richness estimate. Alpha diversity analysis revealed significant variation across intestinal locations, encompassing bacterial (**Figure 2D**), fungal, and viral reads (**Supp. Figure S2B**). Shannon diversity was significantly higher in the cecum and colon compared to the intestine, suggesting a more balanced microbial community structure in the large intestinal regions. The number of observed species and Chao1 richness estimates were also significantly higher in the colon relative to the intestine, but not significantly higher between the cecum and the intestine. No significant differences were detected between the cecum and colon for these metrics. In contrast, fungal and viral communities displayed different spatial trends (**Supp. Figure S2B**). Moreover, region-specific differences were observed in the fungal-to-bacterial and viral-to-bacterial richness ratios, highlighting variation in the relative contribution of low-biomass taxonomic groups across gut regions (**Supp. Figure S2C**). When stratified by location, the small intestine exhibited significantly higher fungal-to-bacterial richness ratios compared to both the colon and the cecum, suggesting a greater relative fungal diversity in this region. A similar pattern was observed for viral richness: viral-to-bacterial richness ratios were significantly higher in the intestine than in the colon and cecum. These findings indicate that the small intestine, while harboring lower total microbial diversity overall, may support proportionally richer communities of fungi and viruses relative to bacteria.

### Top bacterial taxa vary across intestinal regions, revealing regional compositional signatures

Stacked bar plots of the top 30 most abundant taxa revealed distinct patterns of bacterial composition across intestinal regions. Across taxonomic levels (species, genus, and family), bacterial communities in the colon and cecum exhibited greater compositional similarity to each other, whereas the intestinal community displayed a distinctly different profile (**Figure 2E, Supp. Figure S2D**). The top 30 bacterial species, genera, and families account for over 97%, 99%, and 99% of reads, respectively, across all intestinal regions. At the species level, the top 3 taxa that dominated both the colon and cecum were *Phocaeicola vulgatis, Parabacteroides distasonis, and Akkermansia muciniphila*, which are all aerotolerant anaerobes.^31–33^ The intestine was enriched for a different subset of taxa, including *Curtobacterium flaccumfaciens, Salmonella enterica, and Mycobacterium canetti*, which are aerobes, facultative anaerobes, and aerobes, respectively.^34,35^ This trend was consistent at the genus and family levels, where shared dominant taxa between colon and cecum (majorly represented by genera *Phocaeicola, Parabacteroides, and Bacteroides* and families Bacteroidaceae, Tannerellaceae*, and* Akkermansiaceae) contrasted with unique communities in the intestine (majorly represented by genera *Curtobacterium, Salmonella,* and *Mycobacterium* and families Microbacteriaceae, Enterobacteriaceae, Mycobacteriaceae). At the phylum level, the colon and cecum were largely composed of Bacteroidota and Verrucomicrobiota, while the intestine showed a relatively higher contribution from Actinomycetota and Pseudomonadota. The ratio of the Bacteroidota to Bacillota (formerly known as Firmicutes) phyla tends to increase from the intestine to the cecum to the colon, with ratios of 2.86, 17.19, and 41.60, respectively, also in line with other studies.^36^ Dominant taxa at all intestinal regions are summarized in **supplementary tables 2, 3, 4, and 5**.

### Taxonomic and functional gradients define the gut axis

At the species level, several taxa exhibited clear region-specific enrichment patterns (**Figure 2F, Supp. Figure S3**). For example, higher prevalence of *Klebsiella pneumoniae* and *Salmonella enterica* was found in the intestine, while *Bacteroides* and *Alistipes* species showed higher relative abundance in the cecum and colon. Genus-level analysis revealed similar regional stratification. *Enterobacter* and *Ligilactobacillus* were enriched in the intestine, consistent with the more oxygenated environment of the proximal gut. In contrast, *Bacteroides*, *Alistipes*, and *Muribaculaceae* were more abundant in the cecum and colon. Interestingly, *Akkermansia*, a known mucin-degrading bacterium, was also present across all gut segments, with higher relative abundance in the colon. At the family level, Enterobacteriaceae was highly enriched in the intestine, while Bacteroidaceae and Lachnospiraceae were predominant in the cecum and colon. Other families, including Muribaculaceae and Akkermansiaceae, also showed marked enrichment in the distal gut. Across all taxonomic levels, we observed a consistent gradient of microbiota composition along the gut axis, with a shift from facultative anaerobes like Enterobacteriaceae in the small intestine to obligate anaerobes like Bacteroidaceae and Lachnospiraceae in the cecum and colon.

To identify bacterial taxa that may be driving regional differences, we performed Linear Discriminant Analysis Effect Size (LEfSe) analysis (**Figure 2G**). We identified significant location-specific bacterial enrichment patterns across the intestine, cecum, and colon. Genera enriched in the colon and cecum were predominantly associated with anaerobic taxa typical of these gut regions, while taxa enriched in the intestine reflected the oxygenated, bile-exposed, and antimicrobial-rich environment. The LDA bar plots revealed consistent clustering of features by anatomical region, further corroborating the distinct spatial microbial niches along the gastrointestinal tract.

### Taxonomic consistency of gut microbiota across multi-kingdom and mouse-specific **databases.**

To validate the robustness of our taxonomic findings, we compared species, genus, family, and phylum-level results obtained from the Kraken2 PlusPF database with those generated using a mouse gut-specific reference, the Mouse Gastrointestinal Bacteria Catalogue (MGBC) (**Supp. Figure S4A and B)**. At the species level, the cecum and colon communities were highly similar between databases. Both identified *Phocaeicola vulgatus*, *Parabacteroides distasonis*, and *Akkermansia muciniphila* as top taxa in these regions. Additionally, MGBC-unique taxa (e.g., *RC9 MGBC104799*, *Bacteroides acidifaciens*) were not identified in the PlusPF database, reflecting likely differences in the database. MGBC-specific taxa complemented the regional community profiles without contradicting the dominant species identified by PlusPF. In the intestine, both database pipelines highlighted communities dominated by a few genera, including *Curtobacterium* (PlusPF) or *Ligilactobacillus murinus* and uncultured taxa like *UBA3282* (MGBC). At the genus level, patterns again aligned across databases. *Phocaeicola*, *Parabacteroides*, and *Bacteroides* were dominant in the cecum and colon in both analyses, while less characterized genera such as *RC9* and *UBA3282* appeared uniquely in the MGBC output. At the family level, core families like Bacteroidaceae and Tannerellaceae were among the most abundant in the cecum and colon in both databases. Additionally, Lachnospiraceae and Muribaculaceae, well-established mouse-associated families, were recovered with high relative abundance in MGBC, corroborating patterns seen with PlusPF’s Lachnospiraceae and Bacillota-related families. At the phylum level, both databases identified Bacteroidota as dominant in the cecum and colon, with Verrucomicrobiota also ranking highly. Both pipelines showed consistent regional trends, with Bacteroidota and Bacillota/Firmicutes dominating in the large intestine and a greater representation of Actinomycetota/Proteobacteria in the small intestine. Together, these comparisons demonstrate that despite differences in taxonomic resolution and database composition, the major spatial trends in microbial composition were consistent between the comprehensive multi-kingdom PlusPF database and the curated mouse-specific MGBC database. Dominant taxa at all intestinal regions aligned to the MGBC are summarized in **supplementary tables 7, 8, 9, and 10**.

### Functional profiling reveals a shared core and regional specialization

To investigate the spatial organization of microbial functional potential across the gastrointestinal tract, we profiled Gene Ontology (GO) biological processes derived from microbial metagenomic data using HUMAnN3. We visualized the top 30 most abundant GO terms per region using stacked bar plots (**Figure 2E**). Terms that appeared in the top 30 of at least one region were retained, resulting in a total of 41 unique GO terms across all intestinal regions. The small intestine included terms such as rRNA transcription, global gene silencing, and specialized metabolic pathways. The presence of viral life cycle and transporter-associated terms could suggest microbial interactions with both host and phages. The cecum harbored signatures of immune engagement and metabolic flexibility. In contrast, the colon featured the broadest functional repertoire, dominated by carbohydrate degradation and energy harvest pathways such as cellulose catabolism, carbohydrate transport, and SCFA-related processes. Region-specific enrichment patterns were further corroborated by constructing Venn diagrams comparing the top 50 GO terms and all GO terms across locations (**Figure S5A**). These distinct signatures suggest spatially structured microbial roles tailored to each gut niche. For a full list of shared and distinct terms in these regions, please refer to supplementary table 6.

### Region-specific disruption of gut microbial communities by dietary zinc deficiency

To determine how dietary zinc deficiency affects gut microbial composition, mice were fed with either a zinc adequate (ZnA) or a zinc-deficient (ZnD) diet for three weeks (**Figure 3A**). We confirmed that our dietary interventions induced zinc deficiency in the lumen of gastrointestinal tissues throughout the GI tract (**Figure 3A**).

**Figure 3.**
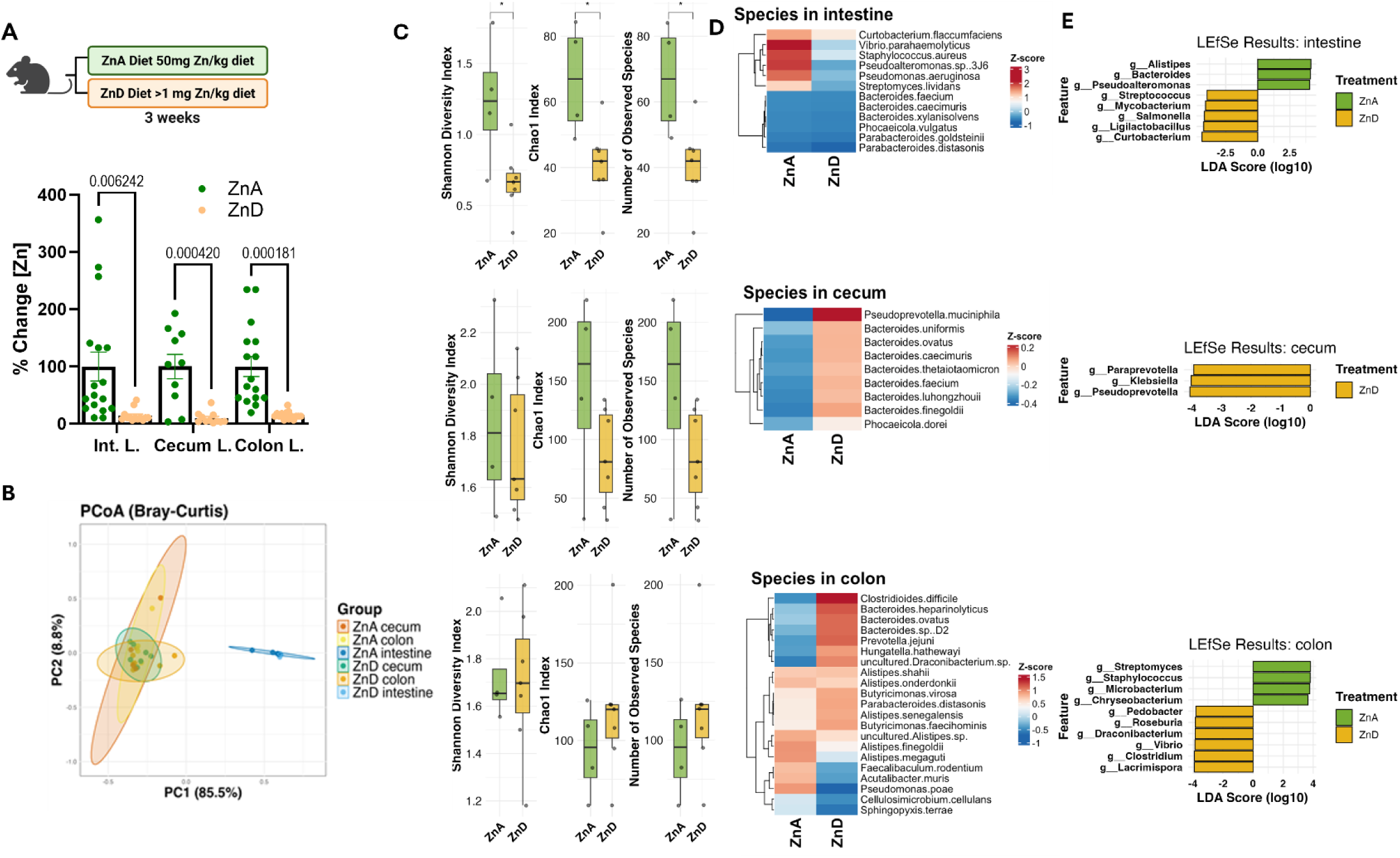
Dietary zinc deficiency drives region-specific dysbiosis and microbial diversity loss in the gut. (A) Luminal zinc quantification. Mice were fed a zinc-adequate (ZnA, 50 mg Zn kg⁻¹) or zinc-deficient (ZnD, <1 mg Zn kg⁻¹) diet for 3 weeks. Luminal contents from the small intestine and colon were digested in trace-metal–grade HNO₃, and Zn concentrations were measured by microwave plasma–atomic-emission spectroscopy (MP-AES; 414.3 nm). Bars show % change ± SE (intestine and colon lumen n = 17 ZnA/17 ZnD; cecum n = 10 ZnA/16 ZnD); points are individual mice. Two-tailed unpaired Student’s t-tests compare diets within each region. (B) Community structure by diet and region. PCoA on Bray–Curtis dissimilarities calculated from log₁₀-CPM-normalized bacterial read counts (Kraken2 + Bracken, PlusPF database) is coloured by treatment and intestinal location. Ellipses indicate 95 % confidence intervals drawn from a multivariate t-distribution. (C) α-diversity effects of zinc deficiency. Shannon diversity, Chao1 richness, and number of observed bacterial species for each intestinal region, separated by treatment (ZnA n = 7, ZnD n = 4 per region). Boxes show the median ± IQR; whiskers extend to 1.5 × IQR; points are samples. Within each region, two-group differences were assessed with Wilcoxon rank-sum tests (* < 0.05, ** < 0.01, *** < 0.001). (D) Region-specific differentially abundant species. Heat-maps display Z-score–scaled relative abundance of bacterial species that were significantly associated with diet in each region (Maaslin2; linear model with diet as fixed effect and cage as random effect, intestine as reference; FDR p < 0.05). Z-scores were calculated across all samples and then averaged within each diet–region stratum. Rows are hierarchically clustered (Euclidean distance, complete linkage). (E) LEfSe biomarkers of zinc status. LEfSe was run separately for each region (CPM normalization, class = Treatment, α(Wilcoxon) = 0.05, log₁₀ LDA cutoff = 1).

To assess how dietary zinc levels impact the gut microbiome, we performed PCoA using Bray-Curtis dissimilarity on samples collected from the small intestine, cecum, and colon of animals fed either a ZnA or ZnD diet (**Figure 3B, Supp. Figure S6A**). When stratified by gut location, distinct clustering of microbial communities based on zinc treatment was pronounced in the intestine, though still evident in the colon and cecum. We measured alpha diversity using the Shannon Index, Chao1 Index, and observed species for bacterial, fungal, and viral communities in the small intestine, cecum, and colon under ZnA and ZnD conditions (**Figure 3C, Supp. Figure S6B**). Bacterial alpha diversity increased from the intestine to the cecum, with the colon displaying intermediate levels. Across all three metrics, the intestine consistently harbored the lowest bacterial diversity, which was further reduced under ZnD conditions. Notably, the reduction in bacterial diversity under ZnD was statistically significant in the intestine (*p* < 0.05), but not in the colon or cecum. In contrast, no significant differences in fungal or viral alpha diversity were observed between ZnA and ZnD conditions at any intestinal location.

To highlight the most prominent taxonomic shifts, we visualized differentially abundant bacterial species and genera identified through comparisons between ZnA and ZnD treatments. Boxplots revealed region-specific differences in bacterial genera between ZnA and ZnD treatments (**Supp. Figure S7A, S7B**). In the intestine, genera such as *Parabacteroides, Phocaeicola, Alistipes, Pseudoalteromonas*, and *Staphylococcus* were present at lower relative abundance in ZnD samples compared to ZnA samples. In the cecum, *Lachnoclostridium* and *Bacteroides* were enriched in ZnA samples, whereas *Pseudoprevotella* was enriched in ZnD samples. The colon exhibited the largest number of differentially expressed taxa, with *Microbacterium*, *Chryseobacterium*, *Pseudomonas*, *Alistipes*, *Butyricimonas*, and *Parabacteroides* enriched in ZnA samples, and *Hungatella* and *Clostridioides* enriched in ZnD samples.

Heatmaps (**Figure 3D, Supp. Fig. S8**) generated using z-scores indicate that several taxa, including *Curtobacterium flaccumfaciens*, *Vibrio parahaemolyticus*, *Staphylococcus aureus*, and *Pseudomonas aeruginosa,* exhibited lower z-score values in ZnD samples compared to ZnA samples, suggesting a trend of reduced abundance. In the cecum, the most pronounced difference was reflected by higher z-score values for *Pseudoprevotella muciniphila* under ZnD conditions, suggesting increased abundance. In the colon, a broader range of bacterial species exhibited varying responses to zinc status. Specifically, *Clostridioides difficile*, *Prevotella jujeni*, and *Hungatella hathewayi* displayed higher z-score values under ZnD conditions, while *Faecalibacterium rodentium* and *Acutalibacter muris* showed higher z-score values under ZnA conditions. Overall, these results highlight the complex, region-specific responses of gut microbiota to zinc status, with the small intestine showing notable shifts towards reduced diversity.

To identify microbial taxa that differentiate between ZnA and ZnD mice within specific intestinal regions, we performed LEfSe analysis on bacterial genus-level profiles from the intestine, cecum, and colon (**Figure 3D**). In the intestine, several genera commonly associated with healthy gut homeostasis, such as *Alistipes, Bacteroides*, and *Pseudoalteromonas*, were identified as having significantly higher LDA scores in ZnA mice, indicating their enrichment under ZnA conditions. In contrast, several genera had lower LDA scores in ZnD samples, reflecting a potential reduction in bacterial diversity. In the cecum, genera enriched under ZnD conditions were identified, including *Paraprevotella*, *Klebsiella*, and *Pseudoprevotella*, as evidenced by their significantly higher LDA scores in ZnD samples. In the colon, genera such as *Streptomyces*, *Staphylococcu*s, *Chryseobacterium*, and *Microbacterium* showed higher LDA scores in ZnA samples, suggesting their enrichment under ZnA conditions. Conversely, ZnD mice exhibited genera with higher LDA scores including *Pedobacter, Roseburia, Draconibacterium, Vibrio, Clostridium*, and *Lacrimospora*. Overall, these findings demonstrate the use of LEfSe analysis for identifying taxa that significantly discriminate between ZnA and ZnD conditions, revealing intestinal region-specific microbial responses to zinc status."

### Zinc status influences microbial functional potential across the gut

To investigate the functional impact of zinc deficiency on gut microbial communities, we identified the top 50 most variable GO terms in the intestine, cecum, and colon and compared their relative abundance between ZnA and ZnD mice (**Supp Figure S9**). In the small intestine, ZnD mice showed increased representation of a few select functions, including DNA repair, ATP hydrolysis, and carbohydrate metabolic processes, but overall exhibited lower relative abundance across most GO terms, indicating a global reduction in microbial metabolic and transcriptional activity under ZnD conditions in this region. In contrast, the cecum and colon showed the opposite trend. The majority of high-variance GO terms in these regions were upregulated under ZnD conditions, including functions related to zinc ion binding, genome modification, metabolism, and signaling. Further suggesting that zinc deficiency could induce a functional remodeling of the microbial community in the large intestine, potentially reflecting stress adaptation or shifts in energy metabolism.

### Conserved taxa exhibit zinc-responsive signatures in mice and humans

To contextualize our findings in a broader, translational framework, we next compared the mouse gut microbiome to publicly available human intestinal metagenomes, focusing on genera that were present in both species and significantly altered by zinc status in our mouse dataset. Comparative taxonomic profiling revealed substantial overlap between human and mouse microbiota at the genus level, with 113 genera shared in the small intestine (**Figure 4A**) and 168 in the colon (**Figure 4C**). Despite broader taxonomic diversity in humans, the overlapping genera comprised key commensals previously linked to gut homeostasis and zinc metabolism, such as *Bacteroides, Alistipes, Parabacteroides*, and *Phocaeicola* in the intestine, while the colon shared genera included *Parabacteroides, Microbacterium, Clostridioides*, and *Pseudomonas*. We next evaluated whether any of these shared genera exhibited significant zinc-driven shifts in our mouse models. Several conserved genera showed significant differential abundance between ZnA and ZnD conditions in mice. In the colon, *Microbacterium*, *Butyricimonas*, *Chryseobacterium*, and *Pseudomonas* were significantly reduced under ZnD conditions, while *Clostridiodes* and *Hungatella* were increased under ZnD conditions (**Figure 4D**). In the small intestine, *Bacteroides*, *Parabacteroides*, *Phocaeicola*, *Pseudoalteromonas*, *Staphylococcus*, and *Streptomyces* were lower in ZnD samples (**Figure 4B**). These patterns indicate that conserved taxa across species are also responsive to zinc status in a region-specific manner.

**Figure 4.**
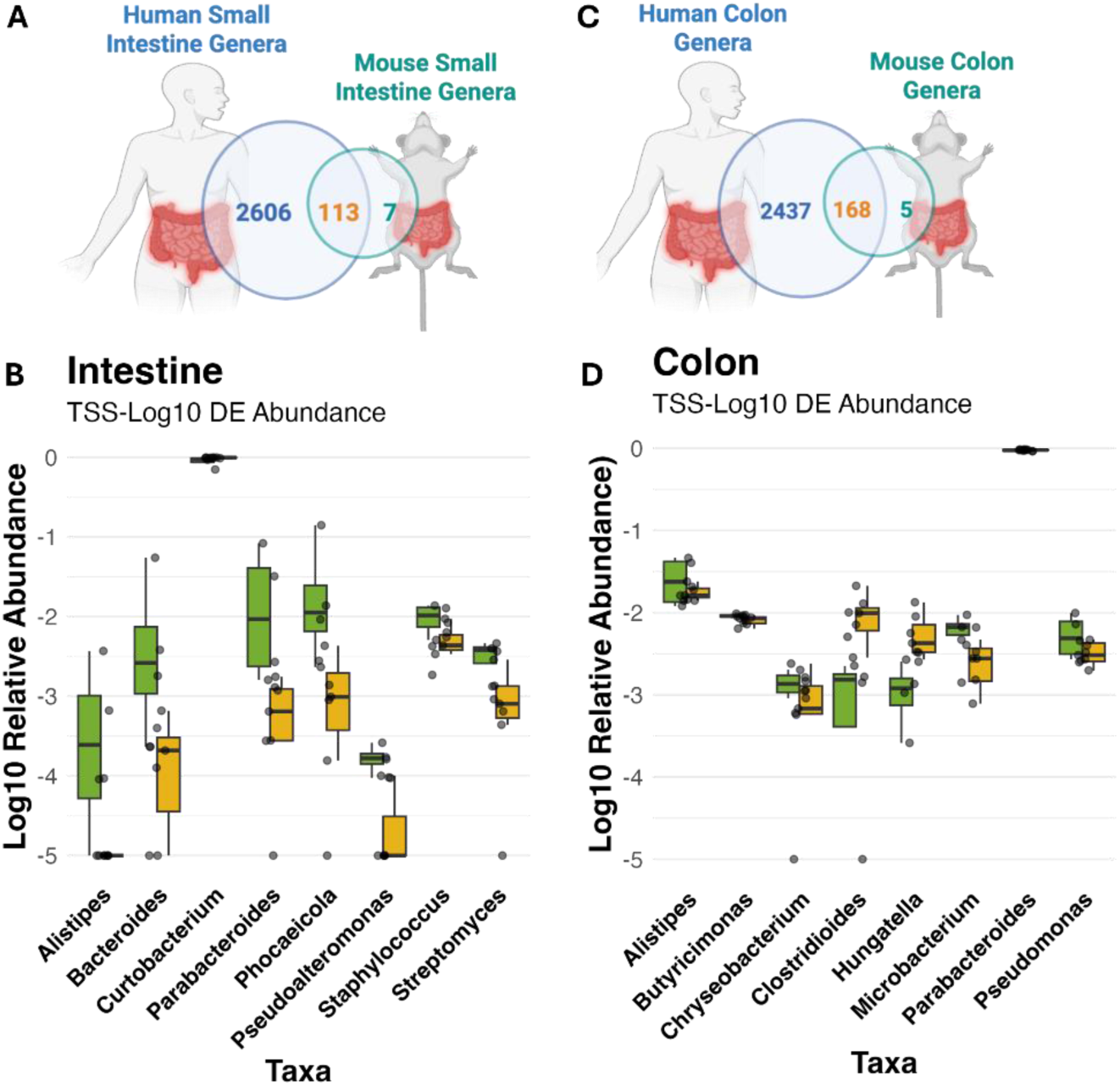
Conserved, zinc-responsive bacterial genera shared between humans and mice. (A, C) Venn diagrams summarizing the genus-level overlap between publicly available human intestinal metagenomes (blue; healthy adults profiled with ingestible capsules² n intestine = 117, n colon = 45) and our mouse samples (n intestine = 11, n colon = 11). Values denote the total number of genera detected in humans, mice, and those shared in the small intestine (A) and colon (C). (B, D) Taxa-specific effects of dietary zinc deficiency on the shared genera. Boxplots show log₁₀-transformed, TSS-normalized relative abundances of genera that occur in both humans and mice, and were significantly different between ZnA and ZnD mice by MaAsLin2 (linear model with Location as fixed effect, FDR < 0.05; intestine used as the reference level). Boxes depict the median ± IQR; whiskers extend to 1.5 × IQR; points are individual mice (ZnA n = 7, ZnD n = 4 per region). Significance between diets was evaluated with two-sided Wilcoxon rank-sum tests (p-values shown when < 0.05).

### Dietary zinc deficiency alters host transcriptome in a regionally dependent fashion

To determine how dietary zinc deficiency affects host intestinal tissues, we conducted RNA sequencing on colon tissue from mice fed a ZnA or ZnD diet. In order to ensure that any ZnD-induced changes in host transcript expression that we detect are relevant and potentially translatable to humans, we compared expressed genes in our ZnA-fedmice and expressed genes from publicly available RNA-seq data from colon mucosal biopsies of human patients.^37^ In both our mouse sequencing data and the available sequencing data, we considered a gene to be expressed if more than 10% of samples displayed at least 10 counts. This analysis revealed 19,521 genes expressed in human colonic mucosa and 10,569 genes in mouse colon tissue. Of these, 10,127 genes were expressed in both human colon mucosa and mouse colon tissue. That is, approximately 96% of the genes expressed in the mouse colon are indeed expressed in humans (**Figure 5A**). Although it has long been known that the human and mouse genomes are largely similar,^38^ our analysis confirms that the local transcriptomes in colon tissue of mice and humans are also highly similar, indicating that mouse models of dietary zinc deficiency are appropriate for study.

**Figure 5.**
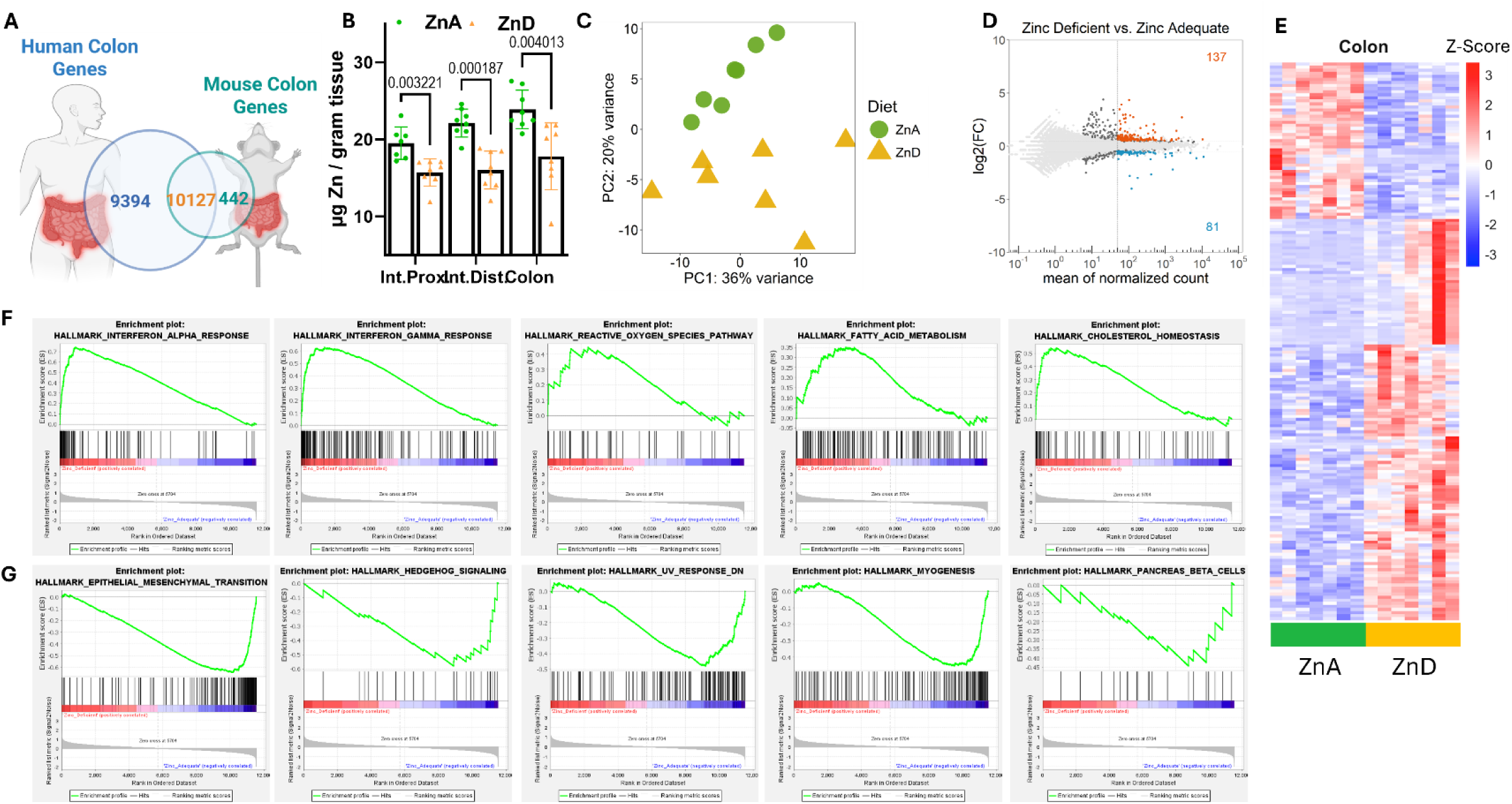
Zinc deficiency alters host gene expression in colon tissue. (A) Overlapping expressed genes in mouse and human colon transcriptomes. (B) Zn concentrations in the proximal small intestine, distal small intestine, and colon tissues of ZnA and ZnD fed mice (n = 6-8). (C) Principal component analysis (PCA) of colon transcriptome of ZnA and ZnD fed mice (n = 6-7). (D) MA plot highlighting differential expressed genes in colon tissues of ZnD mice. Upregulated genes (137) shown in red, downregulated genes (81) in blue. (E) Heatmap of differentially expressed genes (P-adj < 0.05) in colon tissues of ZnD fed mice. Heirarchical clustering was performed using Ward D2 method on genes (rows) and samples (columns). (F, G) Enrichment plots of top 5 upregulated (F) and downregulated (G) gene sets in response to ZnD as determined by GSEA.

Next, we compared the transcriptome of ZnD and ZnA fed mice. We confirmed that our dietary interventions induced a physiologically relevant zinc deficiency in host gastrointestinal tissues throughout the intestinal tract (**Figure 5B**). PCA analysis revealed clear separation between transcriptomes of ZnA- and ZnD-fedmice, suggesting that dietary zinc deficiency induces notable and potentially consequential changes in host gene expression (**Figure 5C**). We conducted differential expression analysis using DESeq2^39^, identifying 218 differentially expressed genes resulting from zinc deficiency. Of these, 137 genes were upregulated and 81 were downregulated (**Figure 5D, E**). To identify what biological functions may be most impacted by zinc deficiency, we conducted gene set enrichment analysis (GSEA).^40,41^ This revealed 17 significantly enriched hallmark gene sets in response to zinc deficiency. Interferon alpha response, interferon gamma response, and reactive oxygen species pathways were the most significantly enriched pathways that were upregulated in ZnD colon tissue (**Figure 5F**). Of those gene sets found to be downregulated in ZnD, epithelial-to-mesenchymal transition, hedgehog signaling, and UV response genes were the most significant (**Figure 5G**). Additionally, GSEA identified *Slc39a4/ Zip4*, as the top-ranked gene in the ranked list of genes used for the analysis, which is appropriate as ZIP4 is known to be the primary transporter responsible for the absorption of dietary zinc in the proximal small intestine. Accordingly, *Zip4* transcript expressions are also sensitive to zinc status. Although the colon has not been considered a significant contributor to zinc absorption, our analysis suggests that *Zip4* may also be responsive to zinc status in the colon. Collectively, our RNA-seq analysis of the impact of dietary zinc deficiency on colon tissues suggests that ZnD promotes inflammatory and oxidative stress signatures while suppressing epithelial cell signaling pathways that are key to maintaining tissue homeostasis. Despite nearly all dietary zinc absorption occurring in the small intestine, the colon also appears to be sensitive to zinc deficiency.

To evaluate the impact of dietary Zn deficiency on the small intestine, we conducted RNA sequencing in proximal small intestine tissue from ZnA and ZnD mice. As before, we first confirmed that approximately 95% of expressed genes in mouse small intestine tissue were expressed in human small intestine tissue (**Figure 6A**). PCA showed clear separation between ZnA and ZnD transcriptomes (**Figure 6B**). In differential expression analysis, we observed a much greater number of differentially expressed genes than in the colon. Of the 1,588 differentially expressed genes, 741 were upregulated and 847 were downregulated (**Figure 6C, D**). GSEA resulted in 17 upregulated 4 downregulated hallmark gene sets in response to Zn deficiency (**Figure 6E**). The most significantly enriched gene sets were similar to those in colon tissue, including the upregulation of genes related to Interferon alpha and gamma responses, and the downregulation of genes involved in the epithelial-to-mesenchymal transition.

**Figure 6.**
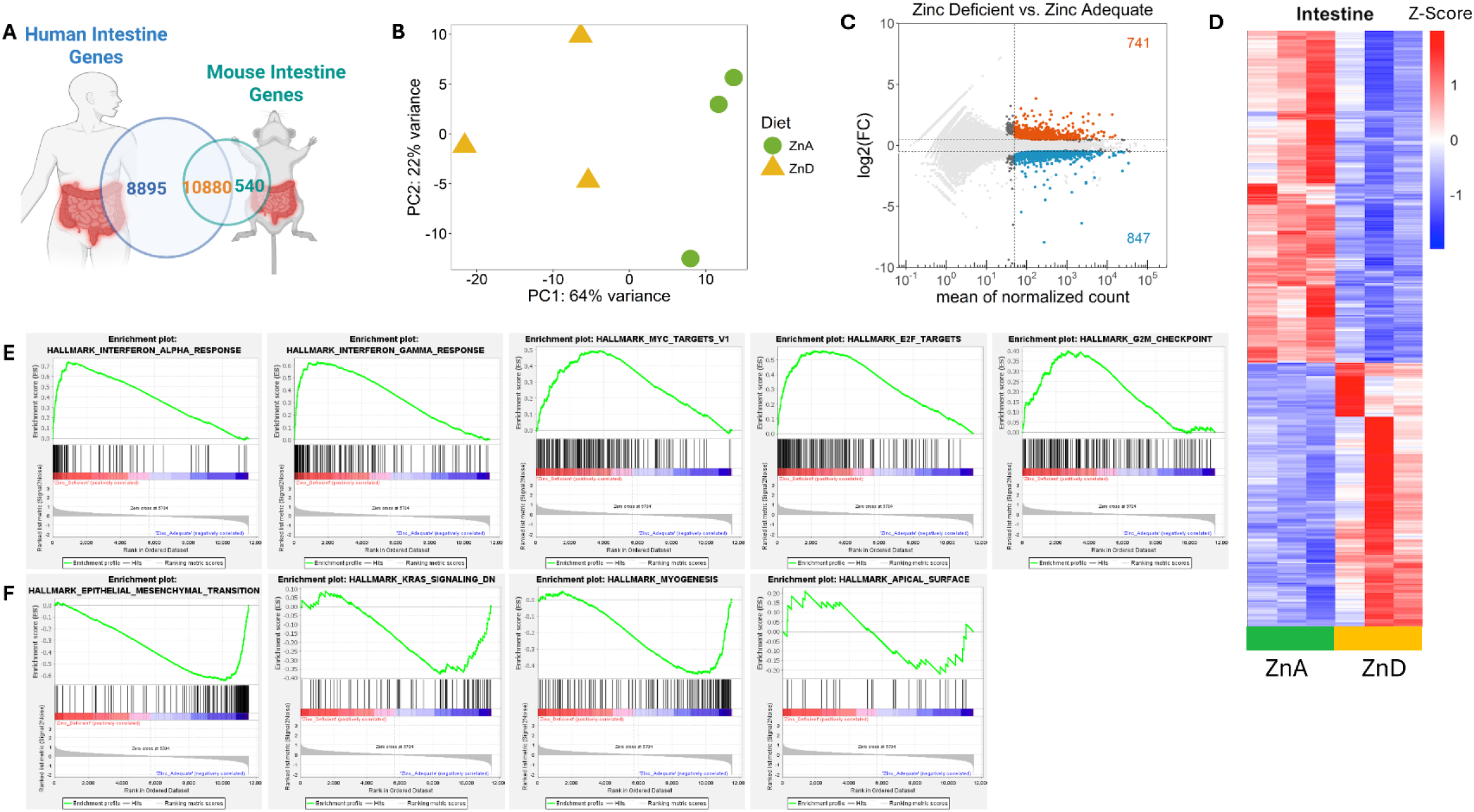
Zinc deficiency alters host gene expression in the proximal small intestine. (A) Overlapping expressed genes in mouse and human small intestine transcriptomes. (B) Principal component analysis (PCA) of the intestine transcriptome of ZnA and ZnD fed mice (n = 3). (C) MA plot highlighting differentially expressed genes in the intestine tissues of ZnD mice. Upregulated genes (741) are shown in red, downregulated genes (847) in blue. (D) Heatmap of differentially expressed genes (P-adj < 0.05) in small intestine tissues of ZnD fed mice. Hierarchical clustering was performed using the Ward D2 method on genes (rows) and samples (columns). (E, F) Enrichment plots of top 5 upregulated (E) and all 4 downregulated (F).

### Zinc deficiency induces region-specific transcriptional alterations and host-microbe associations

Previous findings suggest region-specific differences between the intestine and colon with regard to the significantly enriched genes following dietary zinc deficiency. To elaborate upon these findings, we compared differentially expressed genes in the intestine and colon for shared and tissue-specific regulatory differences (**Figure 7A**). **Figure 7B** shows the top 5 up- and top 5 downregulated GO terms following enrichment analysis for the intestine. Notably, upregulated categories included terms related to xenophagy, intrinsic apoptosis pathways, cellular trafficking, and localized immune system function. Downregulated pathways suggest disruptions in extrinsic apoptosis signaling pathways and impaired immune coordination. **Supplemental Figure 9B** suggests both the intestine and colon share in the upregulation of immune system functions against external stimuli. **Figure 7B** suggests distinct differences in upregulated categories for the colon. Among these are GTPase inhibitor activity, protein localization, and cellular response to hydrogen peroxide, suggesting an increase in regulation of redox pathways, cell migration, and protein targeting. Downregulated pathways include those associated with barrier function and activation of certain immune system functions, which may be a colon-specific compensatory response to the zinc-deficient conditions.

**Figure 7.**
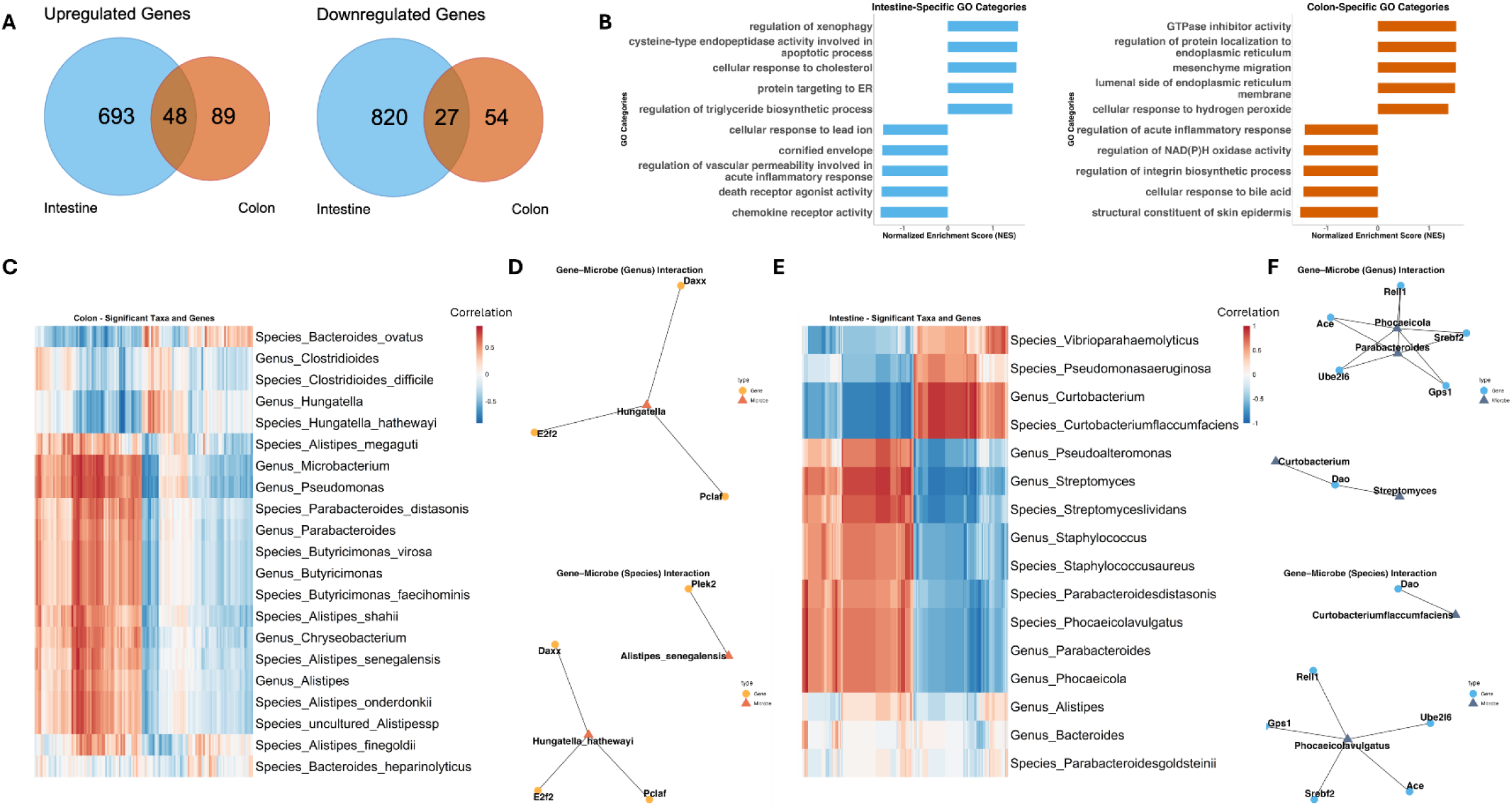
Zinc deficiency results in regionally distinct responses in both host and microbes. (A) Common and unique upregulated and downregulated genes between the small intestine (blue) and colon (red) tissues in ZnD-fed mice. (B) GSEA was performed on uniquely different genes in the colon and small intestine. The top 5 upregulated and downregulated GO categories are shown. (C) Pearson correlation coefficients were calculated between host gene expression and microbial taxa abundance. Significantly different genes unique to the colon tissue and significantly different bacterial genera and species were used. (D) Network visualization of significant gene-microbe correlations (P-adj < 0.2). (E) Pearson correlation coefficients were calculated between host gene expression and microbial taxa abundance. Significantly different genes unique to the small intestine tissue and significantly different bacterial genera and species were used. (F) Network visualization of significant gene-microbe correlations (P-adj < 0.2).

Due to substantial changes in host gene expression and the composition and function of the microbiome throughout the intestinal tract, we sought to profile potential associations between host genes and microbial taxa. To do this, we computed correlations between differentially expressed host genes specific to each tissue **(Figure 7C and E)** and the abundance of significantly different microbial taxa at the genus and species level in both the small intestine and colon. In the colon, we calculated correlations between 143 host genes and 21 microbial taxa, resulting in a total of 3,003 tests (**Figure 7C**). We identified a small network of 6 associations centered around the bacterial species *Hungatella hathewayi,* which was associated with the genes *E2f2, Daxx, and Pclaf* (**Figure 7D**). In the small intestine, we calculated correlations between 1513 host genes and 16 microbial taxa, resulting in 24,208 total tests (**Figure 7E**). This resulted in 17 significant associations between host genes and microbial taxa, with a main network centered around genera *Phocaeicola* and *Parabacteroides*, which were each associated with host genes *Ref1, Ace, Ube2l6, Srebf2*, and *Gps1* (**Figure 7F**). Both taxa were negatively associated with *Ref1, Ace*, and *Ube2l6*, and positively associated with *Srebf2* and *Gps1*. Collectively, our analysis suggests that dietary zinc deficiency has tissue-specific effects on the host transcriptome, leading to regionally distinct interactions between the host and microbes.

## Discussion

This study demonstrates that zinc deficiency induces region-specific alterations in gut microbiota and host-microbe dynamics, revealing important spatial differences in how zinc availability shapes microbial diversity, functional activity, and host gene expression. Using radiotracer and multiomics approaches, our findings uncovered distinct regional microbial compositions and functional adaptations throughout the gastrointestinal tract. Zinc deficiency significantly reduced microbial abundance in the small intestine for all genera. Conversely, the colon exhibited greater resistance to ZnD-driven changes in diversity; however, unlike the small intestine, it showed selective increases in pathogenic bacteria. From the host side, both tissues exhibited upregulation of pathways related to immune function; however, the intestine displayed specific transcriptomic changes in localized apoptosis signals, whereas the colon exhibited reductions in barrier integrity and redox pathways. These results suggest that responses to zinc deficiency are not uniform across the GI tract but are linked to region-specific microbial niches, resulting in region-specific alterations in microbial diversity and transcriptomic regulation. Collectively, these findings provide insights into how zinc deficiency contributes to alterations in host-microbe dynamics and may predispose or exacerbate inflammatory and metabolic diseases.

Studying zinc metabolism and homeostasis in humans is challenging due to its dynamic and complex nature.^42^ Zinc regulatory mechanisms involve intricate processes, including the up- and down-regulation of zinc transport proteins, which are highly responsive to dietary zinc levels and biological signals such as hormones and cytokines.^43–55^ These factors influence the redistribution of zinc within the body, complicating the analysis of its metabolism. Additionally, determining net zinc absorption and loss is challenging when using metal tracers, given the various routes of zinc excretion. To address these limitations, animal models provide a controlled environment for detailed molecular and physiological investigations, offering valuable insights into zinc regulation. The GI tract is characterized by regional differences in structure, function, and luminal conditions across and within tissues. Studying the gut microbiome at a regional level is essential due to differences in pH, oxygen availability, bile acid concentrations, nutrient gradients, mucus composition, and transit time that create unique selective pressures that shape which microbes can colonize and thrive in each region.^56^ This region-specific perspective is particularly relevant for identifying how micronutrient dynamics, such as zinc availability, and local immune environments intersect with microbial communities, influencing both commensal and pathogenic taxa.^57^ Furthermore, profiling of zinc deficiency-driven changes may provide important insights into diseases commonly associated with zinc deficiency, such as IBD, in which previous research has established a critical need for understanding region-specific physiology.^58^

The small intestine is the primary site of dietary zinc absorption. Host enterocyte regulation of zinc uptake by transporters, such as ZIP4 and ZnT1, can create luminal microenvironments of localized zinc scarcity, necessitating a high-affinity zinc uptake system in microbes, which is known to be present in *Salmonella enterica*.^59^ While the relative abundance of this bacterium in the intestine seemingly decreases during zinc deficiency, this finding may be indicative that zinc-binding adaptations may not be sufficient for microbial survival within the small intestine. Furthermore, functional enrichment analysis also suggests that the zinc ion binding is decreased in microbes within the small intestine, yet we see an increase in zinc ion binding pathways in the colon, where we also observed higher microbial diversity compared to the small intestine. Notably, in our transcriptomic analysis, GSEA identified *Slc39a4/Zip4* as the top-ranked gene in the ranked list of genes used for the analysis in both the small intestine and colon. ZIP4 is known to be the primary transporter responsible for the absorption of dietary zinc in the proximal small intestine. Accordingly, *Zip4* transcript expressions are also sensitive to zinc status. Although the colon has not been considered a significant contributor to zinc absorption, our analysis suggests that *Zip4* may also be responsive to zinc status in the colon.

Zinc status also shaped microbial composition, consistent with observations from a study in Zn-deficient broiler chickens, where zinc deficiency led to altered microbial communities and reduced species richness and diversity.^60^ Similarly, in the human study of school-age children, Zn deficiency was associated with higher relative abundances of *Coprobacter*, *Acetivibrio*, *Paraprevotella*, and *Clostridium_XI*.^61^ These findings concur with our ZnD-driven enrichment and depletion of certain gut bacterial groups, such as the increase in *Hungatella, Pseudoprevotella* and *Clostridioides* in the cecum and colon of ZnD samples. As well as depletion of *Parabacteroides, Bacteroides,* and *Alistipes* in the small intestine, and *Lachnoclostridium,* and *Butyricimonas* in cecal and colon ZnD samples. Both studies, in agreement with our own, support the notion that zinc deficiency selectively promotes bacterial taxa with potential zinc-scavenging capabilities, while concurrently reducing taxa associated with beneficial fermentation processes and short-chain fatty acid production.

Enrichment of *Hungatella* in ZnD colon samples mirrors findings in human IBD and colorectal cancer, where *Hungatella hathewayi* was transcriptionally active and pro-inflammatory.^62,63^ Notably, *Hungatella hathewayi* was the main microbial species found to be correlated with host genes that were differentially expressed specifically in colon tissue in response to ZnD, namely *E2f2, Daxx*, and *Pclaf. E2f2* and *Pclaf* have been found to be enriched in mouse models of colitis and patients with acute ulcerative colitis, respectively^64–66^ though potential functional roles for these genes in colitis remain unclear. In addition to IBD, *Hungatella hathewayi* has also been associated with colorectal cancer, and inoculation with this species in mice resulted in an increase in colon epithelial cell proliferation. Each of *Pclaf*, *E2f2*, and *Daxx* is also implicated in colorectal cancer and known to be involved in the regulation of transcription and cell proliferation. Therefore, our findings suggest that *Hungatella hathewayi* is a potential microbial mediator of changes in host transcription and cell proliferation in response to dietary Zn deficiency and may contribute to Zn deficiency as a risk factor for chronic gastrointestinal disease.

*Phocaeicola vulgatus,* formerly *Bacteroides vulgatus,* was found to be decreased in the small intestine of ZnD fed mice, and was the species at the center of the network of gene-microbe associates we identified in the small intestine. This bacterial species has previously been found to be increased in ulcerative colitis patients but has recently been shown to alleviate colitis symptoms in experimental conditions.^67–69^ In our network analysis, *Phocaeicola vulgatus* was found to be associated with *Srebf2*, which was downregulated in the small intestine in response to zinc deficiency, is a key regulator of cholesterol homeostasis, and has been implicated in gastrointestinal inflammation. Abnormalities in lipid metabolism are common in IBD patients, in particular Crohn’s disease.^70,71^ Animals and patients with intestinal injuries were found to have decreased *Srebp2* expression, and *Srebp2*-deficient enterocytes were found to have increased inflammatory responses. We also showed that the increased abundance of *Parabacteroides distasonis* in the small intestine was associated with host genes involved in cellular signaling, lipid metabolism, and protein degradation. Prior studies have linked this bacterium to genes regulating immune function,^72^ supporting our findings that zinc deficiency enhances crosstalk of this bacterium with the host transcriptome, to induce a heightened immune response. Collectively, our studies implicate *Phocaeicola vulgatus and* a *Parabacteroides distasonis* as potential mediators of host-microbe-nutrient interactions that could be relevant for further study in the context of inflammation.

The substantial overlap in genera between mice and humans, particularly in the colon, reinforces the translational value of the mouse model for investigating spatial microbiome dynamics. Importantly, we identified several conserved genera such as *Parabacteroides, Bacteroides*, and *Phocaeicola* that exhibit zinc-responsive abundance changes in the colon and intestine, suggesting these taxa may play conserved roles in zinc sensing or utilization and could serve as potential targets for therapeutic zinc modulation. In the small intestine, Zn-deficient mice showed reduced levels of multiple taxa linked to immune regulation and host health, including *Alistipes*, *Bacteroides*, *Pseudoalteromonas*, and *Streptomyces*.^73–75^ *Parabacteroides*, another core commensal with known anti-inflammatory effects,^76^ was also depleted in the ZnD intestine, supporting the idea that zinc may help preserve beneficial, inflammation-dampening taxa. Intriguingly, *Streptomyces*, a rare but bioactive genus known to produce immunosuppressants and antiproliferatives like rapamycin, was also lost in the ZnD intestine. Its depletion may contribute to the pro-inflammatory environment observed during zinc deficiency and possibly the increased colon cancer risk associated with dysbiosis and chronic inflammation.^77^

Our transcriptomic analysis revealed distinct responses to dietary zinc deficiency in gastrointestinal tissues, with a much larger perturbation in gene expression in the small intestine than the colon. As the primary site of dietary zinc absorption (**Figure 1B**),^21,22^ the small intestine showed nearly 10-fold more differentially expressed genes, yet the colon also displayed significant sensitivity, as shown by clear separation of ZnA and ZnD profiles in clustering analyses (**Figure 5D, E**). Importantly, despite its more modest response, the colon shared key findings with the small intestine, including enrichment of interferon signaling pathways and immune-related functions, as well as regulation of zinc-responsive genes such as the upregulation of Slc39a4/Zip4, a zinc transporter, and downregulation of Mt1 and Mt2, genes involved in zinc binding and storage. While ZIP4 is well known for its role in zinc absorption in the small intestine, ^22^ our findings uniquely highlight its upregulation in the colon during zinc deficiency, suggesting an adaptive mechanism for zinc homeostasis. This novel observation points to the colon’s capacity to potentially compensate for zinc uptake in conditions where the small intestine is impaired, such as Crohn’s Disease.^78^

While both tissues shared alterations in immune response, GSEA analysis of the tissue-specific differentially regulated genes highlighted further distinctions by location. Interestingly, intestine-specific enriched terms pointed to dysregulation in apoptotic signaling pathways. While genes encoding cysteine-specific endopeptidases were upregulated, death receptors promoting apoptosis were downregulated (Figure 7B). While zinc deficiency has been shown to increase caspase-dependent apoptosis,^79^ further understanding of this dysregulation in extrinsic apoptotic signaling is needed. Immune signaling in the intestine was further characterized by increased xenophagy, regulation of cytotoxicity, and ER stress responses, suggesting an active adaptation to cellular stress. In contrast, colon-specific enrichment revealed a pronounced downregulation of pathways crucial for epithelial barrier maintenance and protection against oxidative damage. This finding aligns with previous evidence linking zinc deficiency to elevated oxidative stress.^80^ Notably, oxidative stress-related pathways were enriched only in the colon, highlighting potential region-specific vulnerabilities.

The gastrointestinal tract is composed of multiple layers of specialized tissues with multiple distinct niches. From the outer muscularis externa to the inner mucosa, there exist many different cell types which could experience differential responses to dietary Zn deficiency. Even within the mucosa, resident and recruited immune cells in the lamina propria and absorptive enterocytes in the epithelium may have different responses to dietary Zn deficiency, which is complicated further by potential crosstalk between cell types and tissues. This relationship should be given particular consideration when addressing dietary Zn deficiency as a risk factor for chronic diseases. Disorders such as IBD involve a dysregulation of the host immune response, causing inflammation and potentially leaky gut in the epithelium. Understanding how Zn deficiency impacts each of these components individually and in concert could be important for evaluating the utility of Zn repletion in particular individuals.

## Materials and Methods

### Animal Husbandry and Experimental Conditions

Mice were housed in a standard laboratory rodent facility and maintained on a commercial irradiated chow diet (LabDiet 5053, LabDiet, St. Louis, MO), which contained 79 mg/kg zinc as ZnSO4, with ad libitum access to autoclaved tap water at room temperature (∼23 °C; ∼35% humidity) following a 14:10 hour light dark cycle. Both male and female mice, aged 8–16 weeks, were used as age-matched young adult cohorts across all experimental groups. Euthanasia was performed via exsanguination using cardiac puncture under isoflurane anesthesia. Injections and oral gavage procedures were administered under anesthesia to minimize animal distress. All experimental protocols complied with the standards approved by the Cornell University Institutional Animal Care and Use Committee (IACUC). *Dietary Zinc Deficiency:* Mice were fed with either zinc adequate [ZnA, 50 mg Zn/kg diet-Teklad, Inotiv, IN, USA (TD 85420)] or low zinc [ZnD, <1mg Zn/kg diet-Teklad, Inotiv, IN, USA (TD 85419)] for three weeks in shoebox cages.^81^

*Fecal transplant:* Fecal pellets were collected from in-house C57BL6 mice maintained in a specific-pathogen-free (SPF) facility to prepare the fecal slurry. Germ-Free, C57BL/6 mice (Taconic Biosciences, Cat # GF-B6) were inoculated by oral gavage for three consecutive days with 0.2 ml of the fecal slurry.^81–83^ Following conventionalization, mice were maintained in sterile cages for three weeks. At the end of the three weeks, blood and tissue were collected for tissue metal analysis.

### Tissue Metal Quantification

Metal concentrations in tissues were determined using microwave plasma–atomic emission spectrometry (MP-AES; Agilent Technologies, Santa Clara, CA) following overnight digestion at 80°C and subsequent dilution in deionized water tailored to the tissue type. ^30–33^ Tissue metal levels were normalized based on wet tissue weight. Blood samples were collected via cardiac puncture into EDTA-coated tubes, and plasma was harvested by centrifugation at 3000 × g for 15 minutes. ^34^ Plasma samples were diluted 1:5 in deionized water prior to MP-AES analysis.

### 65Zn Transport Assay

To evaluate zinc transport, mice received ^65^Zn isotope (Eckert & Ziegler, Valencia, CA) either by oral gavage (4 μCi/mouse in 100 μL) to measure mucosal to serosal Zn transport, or subcutaneous injection (2 μCi/mouse in 100 μL) to examine serosal to mucosal transport. Tissues were harvested 3 hours post-administration, as described previously. ^35–37^ Intestinal tissues were perfused with a metal-chelating buffer containing 10 mM EDTA, 10 mM HEPES, and 0.9% NaCl to remove extracellularly bound metals. Radioactivity in whole tissues was measured using a gamma counter and normalized to wet tissue weight.

### Luminal Collection and Storage

Following dissection, the colon and small intestine were separated, and luminal contents from the respective sections were flushed with sterile PBS using a 3 mL syringe and a disposable gavage needle. The luminal flush was centrifuged at 6100 xg for 10 minutes at 4°C to pellet digesta containing bacterial and fungal cells. The luminal supernatant and the digesta were resuspended in 100 µL of DNA/RNA Shield (Zymo Cat# R1100-50) and stored at -80°C prior to DNA extraction. The cecal contents were also collected by puncturing the cecum and storing them at -80°C prior to DNA extraction.

### DNA Extraction and metagenomics sequencing

Intestinal and colon digesta and frozen cecal samples were extracted using the Qiagen QIAmp Fast DNA Stool Mini Kit (Qiagen, Cat #51604), with 180–200 mg of stool used as input material per extraction. The DNA was eluted in Buffer AE (Qiagen, Cat# 51604) and stored at −20 °C before further processing. DNA concentrations were measured using a NanoDrop spectrophotometer (Thermo Fisher Scientific, Cat# ND-2000). Library preparation was performed with the Watchmaker Genomics Library Prep Kit (Watchmaker Genomics, Cat# 7K0022-096), incorporating Truseq-compatible IDT xGen UDI-UMI adapters to fragmented metagenomic DNA (IDT, Cat# 10005903). DNA libraries were quantified using qPCR with primers targeting the P5 and P7 Illumina adapter sequences (P7: CAA GCA GAA GAC GGC ATA CGA) and P5 primer (P5:AAT GAT ACG GCG ACC ACC GA). Amplification was performed using SYBR Green Master Mix (Applied Biosystems, Thermo Fisher Cat# A46109) on the QuantStudio 3 instrument. Libraries were normalized to 10 nM and quantified again using qPCR amplified using P5/P7 universal adapters. Normalized libraries were pooled at a final concentration of 1nM prior to sequencing. Sequencing was conducted on the Illumina NextSeq 500 platform at the Cornell University Institute of Biotechnology.

### Quality control for metagenomics sequencing data

Raw sequencing data was received in FASTQ format and processed on the Cornell BioHPC platform, a cloud-based computational environment using the Linux operating system. Quality control of the FASTQ files was performed using FastQC (v0.12.1) (https://www.bioinformatics.babraham.ac.uk/projects/fastqc/), and summary reports were aggregated using MultiQC (v1.15). To ensure comparability in sequencing depth across samples within intestinal regions and dietary treatment groups, we examined the distribution of total reads per sample. Samples with disproportionately higher sequencing depth were randomly downsampled to 1 million reads each using seqtk (v1.2-r102-dirty).^84^

### Aligning Metagenomic Reads/Taxonomic Assignment

Metagenomic sequencing reads were processed for taxonomic classification using Kraken2 (v2.1.3). Reads were aligned against two reference databases to capture both broad microbial diversity and mouse gut-specific bacterial profiles. First, all samples were aligned to the Kraken2 PlusPF database, a comprehensive database comprising reference sequences from bacterial, archaeal, viral, protozoan, and fungal genomes, as well as plasmids, and the human genome (GRCh38). To remove host-derived sequences, raw reads from the small intestine were aligned to the *Mus musculus* reference genome GRCm39 using Bowtie v1.1.2, and only unmapped reads were retained for downstream functional profiling. To enhance taxonomic resolution specific to the mouse gut microbiome, we also aligned reads to the MGBC Mouse Gut Database, a curated database derived from the Mouse Gastrointestinal Bacteria Catalogue (MGBC), containing 26,640 high-quality bacterial genomes representative of mouse gut-associated bacteria. This database allows for the classification of species and strains commonly found in the murine intestinal environment. For both databases, Bracken (v2.9) was used to re-estimate taxonomic abundances from Kraken2 classification reports. Bracken analyses were conducted at the species, genus, family, and phylum levels for both databases. Additionally, kingdom-level classification was performed for the PlusPF database only, given its inclusion of fungal and viral sequences.

### Taxonomic Profiling and Visualization by Intestinal Location

To evaluate the distribution of bacterial communities across different regions of the mouse gastrointestinal tract, we generated stacked bar charts stratified by taxonomic rank and sample location. The top 30 most abundant taxa at each level were retained based on their cumulative relative abundance. Average abundances were then calculated for each intestinal location and visualized using ggplot2.

### Beta Diversity by Intestinal Regions and Treatment

To assess spatial and treatment-driven variation in microbial community composition, we performed beta diversity analyses separately for bacterial, fungal, and viral taxonomic profiles, as well as a combined analysis including all reads. Principal Coordinates Analysis (PCoA) was conducted for each domain and for the combined dataset using Bray-Curtis dissimilarity. To normalize for differences in library size, relative abundance was calculated from Bracken-derived reads for each sample by dividing feature counts by the total counts per sample. Bray-Curtis distances were computed from the normalized matrices, followed by classical multidimensional scaling. The first two principal coordinates were extracted for visualization. Ellipses representing the 95% confidence intervals were added.

### Alpha Diversity and Inter-Kingdom Ratio Analyses of the Gut Microbiome by Intestinal Region and Zinc Status

Raw read counts were classified at the kingdom level using Kraken2 and Bracken, resulting in separate read count data for bacteria, fungi, and viruses. To calculate relative abundance, counts for each kingdom were divided by the total reads per sample.

To assess the alpha diversity of the gut microbiome across intestinal regions, we computed Shannon diversity, Chao1 richness, and observed species metrics using the the phyloseq package in R. Analyses were performed on both aggregate microbial communities (bacteria, fungi, and viruses combined) and kingdom-specific datasets using the PlusPF database. Diversity metrics were calculated from Bracken-derived count matrices normalized by total sum scaling.

The wilcox.test used to assess statistical significance between groups. Pairwise comparisons were visualized using boxplots, and statistically significant comparisons were annotated.

To compare the relative richness of bacterial, fungal, and viral communities across treatment groups and intestinal locations, we computed domain-specific species richness as the number of observed taxawith nonzero read counts per sample. For each sample, we calculated two richness ratios: (1) the fungal-to-bacterial richness ratio, and (2) the viral-to-bacterial richness ratio. Pairwise Wilcoxon rank-sum tests were used to assess significance between groups.

### Taxonomic Profiling of Gut Microbiota Across Multi-Kingdom and Mouse-Databases

To assess the robustness and reproducibility of our taxonomic classifications, we performed comparative profiling of gut microbiota using two distinct Kraken2-compatible databases: the comprehensive PlusPF database and the Mouse Gastrointestinal Bacteria Catalogue (MGBC). The PlusPF database was used for primary analyses due to its broad coverage of bacterial, fungal, and viral taxa, enabling multi-kingdom characterization of the metagenomic dataset. To validate these findings in a host-relevant context, the same raw sequence data was analyzed using the MGBC database. Species, genus, family, and phylum level taxonomic profiles were generated from both pipelines and compared across intestinal regions. Key shared and unique taxa between databases were identified at each taxonomic level, and relative abundance patterns were evaluated to determine consistency across database sources.

### Functional Profiling

Metagenomic functional profiling was performed using HUMAnN3 (v3.7), using default parameters for mapping against the ChocoPhlAn and UniRef90 databases. For each sample, gene family abundance tables (genefamilies.tsv) were normalized to copies per million (CPM) using humann_renorm_table. To facilitate pathway- and ontology-level analyses, gene families were regrouped to Gene Ontology (GO) terms using humann_regroup_table with the uniref90_go mapping option. Normalized and regrouped tables were joined across all samples using humann_join_tables to create a unified GO abundance matrix. The only group of samples that had a significant amount of mouse host reads was the subset of small intestinal samples. Mouse reads were removed from the intestinal samples prior to running HUMAnN3 on those samples. GO term abundance tables (in counts per million, CPM) were imported and preprocessed in R using the AnnotationDbi and GO.db R packages. To identify the most variable functions across intestinal locations, we calculated the row-wise variance of GO term abundances. A total of 2802 unique Gene Ontology (GO) terms were detected across the metagenomic dataset (small intestine = 840; cecum = 2343, colon = 2607). Applying an abundance-prevalence filter (≥ 10 counts per million in at least one region and presence in ≥ 2 regions) removed low-confidence and single-sample noise, but preserved the overall regional pattern. The high-confidence set retained 748 terms in the small intestine, 1678 terms in the cecum, and 1679 terms in the colon. We visualized overlaps between the top 50 GO terms across regions using Venn diagrams and categorized terms as unique to a single region, shared between two regions, or conserved across all three. Additionally, a stacked bar chart of the top 30 most abundant GO terms per region was generated to highlight dominant functional profiles in each anatomical niche.

### Linear Discriminant Analysis

Linear Discriminant Analysis Effect Size (LEfSe) analysis was performed using the microbiomeMarker R package (v1.2.1) for bacterial reads to identify differentially abundant taxa by treatment and by location. Count normalization was performed using the counts per million (CPM) method, with an LDA cutoff of 1 applied for treatment comparisons within each intestinal region. A more stringent cutoff of 1.5 was used for comparisons between locations when samples were not stratified by treatment. For each location, the data was subset to include only samples from that region, and remove taxa with zero variance within either treatment group.

### Heatmap of Functionally Annotated Metagenomes Using Humann3

To visualize spatial differences in microbial functional profiles between zinc-adequate (ZnA) and zinc-deficient (ZnD) conditions, we generated location-specific heatmaps of Gene Ontology (GO) terms. This analysis focused on the top 50 most variable GO terms per anatomical region based on metagenomic functional abundance data obtained from HUMAnN3 and annotated with GO terms. GO terms were filtered to include only those that had non-zero abundance in both ZnA and ZnD treatment groups and the top 50 most variable GO terms were selected. For each GO term, mean relative abundance was calculated separately for ZnA and ZnD groups, producing a two-column matrix per region. Heatmaps were row-scaled to emphasize relative shifts in abundance between treatments. Hierarchical clustering was applied to rows using Euclidean distance.

### Differential Abundance Analysis Using MaAsLin2

Differential abundance of microbial taxa across treatment groups and intestinal locations was assessed using MaAsLin2 (Multivariate Association with Linear Models). Analyses were conducted independently for each intestinal region at four taxonomic levels: species, genus, family, and phylum. Taxa present in fewer than 20% of samples were excluded to reduce sparsity and improve statistical power. Bracken-derived abundance tables were used as input, normalized via Total Sum Scaling (TSS), and log-transformed with a small pseudocount (+1e-5) to mitigate skewness. Taxa with a p-value < 0.05 were retained for visualization. Z-score normalization was applied across taxa to enable relative comparison of abundance patterns by location or by treatment. Z-scores were then aggregated by computing the mean per taxon across samples within each location or treatment group prior to visualization on a heatmap. To generate boxplots that captured relative differences in abundance by treatment, additional filtering was applied to ensure that taxa were present in both ZnA and ZnD groups within a given region. In contrast, heatmaps were not subjected to this filter, allowing for visualization of both differential abundance and presence/absence patterns to note taxa that may be uniquely enriched or depleted under zinc-deficient conditions.

### Overlap of Human and Mouse Genera in the Intestine and Colon

To assess taxonomic overlap and zinc-responsive microbial dynamics between human and mouse intestinal microbiomes, we leveraged metagenomic data from the from Shalon et al. (2023).^29^ We downloaded publicly available fastq files corresponding to intestinal capsules 2 and 3 representing the small intestine and capsule 4 representing the ascending colon. These human samples were processed using the same pipeline applied to our mouse metagenomic data: quality-filtered reads were classified taxonomically using Kraken2 and Bracken aligned to the PlusPF database. Bracken outputs were aggregated at species, genus, family, phylum, and kingdom levels. Taxonomic profiles were compared across matched intestinal regions (small intestine and colon) between species. Shared taxa were identified by intersecting genus-level profiles between humans and mice. To examine functional relevance, we extracted the subset of shared genera that were also significantly differentially abundant in our ZnD vs. ZnA mouse data based on Maaslin2 differential abundance analysis. These zinc-responsive, cross-species taxa were visualized using TSS-normalized, log10-scaled boxplots for each region.

### RNA-sequencing

Libraries were generated from isolated RNA with Lexagen QuantSeq 3’ mRNA-Seq Library Prep Kit FWD (Lexogen, Cat# 192.96). Libraries were sequenced on an Illumina NextSeq500 instrument using 75 bp paired-end reads. Reads were quantified with Salmon using Gencode v38 as the reference genome. Differentially expressed genes were identified using DESeq2 with the Benjamini-Hochberg false discovery rate (FDR) correction at a threshold of 0.05. Gene ontology was conducted from significantly different genes using ToppGene ^38^.

### Statistical Analysis

Raw data collection, analysis, and quantification were performed using Microsoft Excel. Graphs and statistical analyses were conducted with GraphPad Prism versions 7–10. For comparisons between groups, statistical significance was assessed using unpaired, two-tailed Student’s t-tests or one-way ANOVA. Data points are represented as means, with error bars indicating ± SEM. A threshold of P < 0.05 was considered statistically significant across all experiments. Detailed statistical parameters and the number of biological replicates used for each experiment are provided in the figure legends. Shotgun metagenomics and RNA sequencing data analysis, including statistical testing, pathway analysis, and graph generation, was performed using RStudio versions 4.4 and 4.2. Figures and experimental designs were created in PowerPoint, with select illustrations sourced from BioRender (https://www.biorender.com/).

## Data Availability

The authors confirm that the data supporting the findings of this study are available within the article [and/or] its supplementary materials. The RNA-seq and metagenomic sequencing reads generated in this study are available in the NCBI Sequence Read Archive (SRA) under BioProject accession numbers PRJNA1285101 and PRJNA1279680.

## Author Contributions

TBA conceived and designed research; TBA, SS, SBM, AAG, and LT performed experiments; TBA, SS, SBM, AAG, and TLT analyzed data; TBA, SS, SBM, AAG interpreted results of experiments; TBA, SS, SBM, AAG prepared figures; TBA, SS, SBM, AAG, and TLT drafted the manuscript; TBA, SS, SBM, AAG, and TLT edited and revised the manuscript; TBA, SS, SBM, AAG, TLT approved the final version of the manuscript.

## Acknowledgments

Shotgun metagenome and RNA sequencing were conducted by the Biotechnology Resource Center (BRC) Genomics Facility (RRID: SCR 021727) at the Cornell Institute of Biotechnology.

## References

1. Chen L, Wang Z, Wang P, et al. Effect of Long-Term and Short-Term Imbalanced Zn Manipulation on Gut Microbiota and Screening for Microbial Markers Sensitive to Zinc Status. Microbiol Spectr. 2021;9(3). doi:10.1128/SPECTRUM.00483-21

2. Popovic A, Bourdon C, Wang PW, et al. Micronutrient supplements can promote disruptive protozoan and fungal communities in the developing infant gut. Nature Communications 2021 12:1. 2021;12(1):1–13. doi:10.1038/s41467-021-27010-3

3. Cheng J, Kolba N, Tako E. The effect of dietary zinc and zinc physiological status on the composition of the gut microbiome in vivo. Crit Rev Food Sci Nutr. Published online 2023. doi:10.1080/10408398.2023.2169857

4. Gordon SR, Vaishnava S. Zinc supplementation modulates T helper 17 cells via the gut microbiome. The Journal of Immunology. 2019;202(1_Supplement):191.13–191.13. doi:10.4049/JIMMUNOL.202.SUPP.191.13

5. Olechnowicz J, Tinkov A, Skalny A, Suliburska J. Zinc status is associated with inflammation, oxidative stress, lipid, and glucose metabolism. Journal of Physiological Sciences. 2018;68(1):19–31. doi:10.1007/S12576-017-0571-7,

6. Fernández-Cao JC, Warthon-Medina M, Moran VH, et al. Zinc Intake and Status and Risk of Type 2 Diabetes Mellitus: A Systematic Review and Meta-Analysis. Nutrients. 2019;11(5):1027. doi:10.3390/NU11051027

7. Zupo R, Sila A, Castellana F, et al. Prevalence of Zinc Deficiency in Inflammatory Bowel Disease: A Systematic Review and Meta-Analysis. Nutrients 2022, Vol 14, Page 4052. 2022;14(19):4052. doi:10.3390/NU14194052

8. WHO. The World Health Organization Report 2002: reducing risks, promoting healthy life. WHO Library Cataloguing-in Publication Data. Published online 2002:232.

9. Ananthakrishnan AN, Khalili H, Song M, Higuchi LM, Richter JM, Chan AT. Zinc intake and risk of Crohn’s disease and ulcerative colitis: a prospective cohort study. Int J Epidemiol. 2015;44(6):1995–2005. doi:10.1093/IJE/DYV301

10. Peng X, Yang Y, Rao Zhong ·, et al. Zinc and Inflammatory Bowel Disease: From Clinical Study to Animal Experiment. Biological Trace Element Research 2024. Published online May 28, 2024:1–11. doi:10.1007/S12011-024-04193-6

11. Sakurai K, Furukawa S, Katsurada T, et al. Effectiveness of administering zinc acetate hydrate to patients with inflammatory bowel disease and zinc deficiency: a retrospective observational two-center study. Intest Res. 2021;20(1):78–89. doi:10.5217/IR.2020.00124

12. Soares NRM, de Moura MSB, de Pinho FA, et al. Zinc supplementation reduces inflammation in ulcerative colitis patients by downregulating gene expression of Zn metalloproteins. PharmaNutrition. 2018;6(3):119–124. doi:10.1016/J.PHANU.2018.06.004

13. Daneshvar M, Ghaheri M, Safarzadeh D, et al. Effect of zinc supplementation on glycemic biomarkers: an umbrella of interventional meta-analyses. Diabetol Metab Syndr. 2024;16(1). doi:10.1186/S13098-024-01366-0

14. Pompano LM, Boy E. Effects of Dose and Duration of Zinc Interventions on Risk Factors for Type 2 Diabetes and Cardiovascular Disease: A Systematic Review and Meta-Analysis. Advances in Nutrition. 2020;12(1):141. doi:10.1093/ADVANCES/NMAA087

15. Wang X, Wu W, Zheng W, et al. Zinc supplementation improves glycemic control for diabetes prevention and management: a systematic review and meta-analysis of randomized controlled trials. Am J Clin Nutr. 2019;110(1):76–90. doi:10.1093/AJCN/NQZ041

16. Yang HY, Hung KC, Chuang MH, et al. Effect of zinc supplementation on blood sugar control in the overweight and obese population: A systematic review and meta-analysis of randomized controlled trials. Obes Res Clin Pract. 2023;17(4):308–317. doi:10.1016/J.ORCP.2023.06.003

17. Zargar AH, Shah NA, Masoodi SR, et al. Copper, zinc, and magnesium levels in non-insulin dependent diabetes mellitus. Postgrad Med J. 1998;74(877):665–668. doi:10.1136/PGMJ.74.877.665

18. Attia JR, Holliday E, Weaver N, et al. The effect of zinc supplementation on glucose homeostasis: a randomised double-blind placebo-controlled trial. Acta Diabetol. 2022;59(7):965. doi:10.1007/S00592-022-01888-X

19. El Dib R, Gameiro OL, Ogata MS, et al. Zinc supplementation for the prevention of type 2 diabetes mellitus in adults with insulin resistance. Cochrane Database Syst Rev. 2015;2015(5). doi:10.1002/14651858.CD005525.PUB3

20. Marriott BP., Marriott BP /Birt, DF., Stallings VA. Present Knowledge in Nutrition. Published online 2020.

21. Lee HH, Prasad AS, Brewer GJ, Owyang C. Zinc absorption in human small intestine. Am J Physiol Gastrointest Liver Physiol. 1989;256(1). doi:10.1152/AJPGI.1989.256.1.G87

22. Ohashi W, Hara T, Takagishi T, Hase K, Fukada T. Maintenance of Intestinal Epithelial Homeostasis by Zinc Transporters. Digestive Diseases and Sciences 2019 64:9. 2019;64(9):2404–2415. doi:10.1007/S10620-019-05561-2

23. Krebs NF, Hambidge KM. Zinc metabolism and homeostasis: The application of tracer techniques to human zinc physiology. BioMetals. 2001;14(3-4):397–412. doi:10.1023/A:1012942409274/METRICS

24. Marriott BP, Birt DF, Stallings MS VA, Yates MSPH RD AA. Present Knowledge in Nutrition: Basic Nutrition and Metabolism.; 2020.

25. Kastl AJ, Terry NA, Wu GD, Albenberg LG. The Structure and Function of the Human Small Intestinal Microbiota: Current Understanding and Future Directions. CMGH. 2020;9(1):33–45. doi:10.1016/j.jcmgh.2019.07.006

26. Kastl AJ, Terry NA, Wu GD, Albenberg LG. The Structure and Function of the Human Small Intestinal Microbiota: Current Understanding and Future Directions. CMGH. 2020;9(1):33–45. doi:10.1016/j.jcmgh.2019.07.006

27. Lkhagva E, Chung HJ, Hong J, et al. The regional diversity of gut microbiome along the GI tract of male C57BL/6 mice. BMC Microbiol. 2021;21(1). doi:10.1186/S12866-021-02099-0,

28. Folz J, Culver RN, Morales JM, et al. Human metabolome variation along the upper intestinal tract. Nature Metabolism 2023 5:5. 2023;5(5):777–788. doi:10.1038/s42255-023-00777-z

29. Shalon D, Culver RN, Grembi JA, et al. Profiling the human intestinal environment under physiological conditions. Nature. 2023;617(7961):581–591. doi:10.1038/S41586-023-05989-7,

30. Wood DE, Lu J, Langmead B. Improved metagenomic analysis with Kraken 2. Genome Biol. 2019;20(1):1–13. doi:10.1186/S13059-019-1891-0/FIGURES/2

31. Ouwerkerk JP, van der Ark KCH, Davids M, et al. Adaptation of Akkermansia muciniphila to the oxic-anoxic interface of the mucus layer. Appl Environ Microbiol. 2016;82(23):6983–6993. doi:10.1128/AEM.01641-16,

32. Baughn AD, Malamy MH. The strict anaerobe Bacteroides fragilis grows in and benefits from nanomolar concentrations of oxygen. Nature. 2004;427(6973):441–444. doi:10.1038/NATURE02285,

33. Ezeji JC, Sarikonda DK, Hopperton A, et al. Parabacteroides distasonis: intriguing aerotolerant gut anaerobe with emerging antimicrobial resistance and pathogenic and probiotic roles in human health. Gut Microbes. 2021;13(1). doi:10.1080/19490976.2021.1922241,

34. Andino A, Hanning I. Salmonella enterica: Survival, colonization, and virulence differences among serovars. Scientific World Journal. 2015;2015. doi:10.1155/2015/520179,

35. Tegli S, Cerboneschi M, Vidaver AK. CHAPTER 12: Detection of Curtobacterium flaccumfaciens pv. flaccumfaciens in Bean Seeds and in Seeds of Other Leguminosae Crops. *Detection of Plant-Pathogenic Bacteria in Seed and Other Planting Material*, Second Edition. Published online January 2017:77–83. doi:10.1094/9780890545416.012

36. McCallum G, Tropini C. The gut microbiota and its biogeography. Nat Rev Microbiol. 2024;22(2):105–118. doi:10.1038/S41579-023-00969-0,

37. Argmann C, Hou R, Ungaro RC, et al. Biopsy and blood-based molecular biomarker of inflammation in IBD. Gut. 2023;72(7):1271–1287. doi:10.1136/GUTJNL-2021-326451

38. Waterston RH, Lindblad-Toh K, Birney E, et al. Initial sequencing and comparative analysis of the mouse genome. Nature. 2002;420(6915):520–562. doi:10.1038/NATURE01262;KWRD=SCIENCE

39. Love MI, Huber W, Anders S. Moderated estimation of fold change and dispersion for RNA-seq data with DESeq2. Genome Biol. 2014;15(12):1–21. doi:10.1186/S13059-014-0550-8/FIGURES/9

40. Mootha VK, Lindgren CM, Eriksson KF, et al. PGC-1α-responsive genes involved in oxidative phosphorylation are coordinately downregulated in human diabetes. Nat Genet. 2003;34(3):267–273. doi:10.1038/NG1180;KWRD=BIOMEDICINE

41. Subramanian A, Tamayo P, Mootha VK, et al. Gene set enrichment analysis: A knowledge-based approach for interpreting genome-wide expression profiles. Proc Natl Acad Sci U S A. 2005;102(43):15545–15550. doi:10.1073/PNAS.0506580102/SUPPL_FILE/06580FIG7.JPG

42. Hambidge KM, Miller L V., Westcott JE, Sheng X, Krebs NF. Zinc bioavailability and homeostasis. American Journal of Clinical Nutrition. 2010;91(5). doi:10.3945/AJCN.2010.28674I,

43. Besecker B, Bao S, Bohacova B, Papp A, Sadee W, Knoell DL. The human zinc transporter SLC39A8 (Zip8) is critical in zinc-mediated cytoprotection in lung epithelia. Am J Physiol Lung Cell Mol Physiol. 2008;294(6). doi:10.1152/AJPLUNG.00057.2008,

44. Merriman C, Fu D. Down-regulation of the islet-specific zinc transporter-8 (ZnT8) protects human insulinoma cells against inflammatory stress. Journal of Biological Chemistry. 2019;294(45):16992–17006. doi:10.1074/jbc.RA119.010937

45. Woodruff G, Bouwkamp CG, De Vrij FM, et al. The zinc transporter SLC39A7 (ZIP7) is essential for regulation of cytosolic zinc levels s. Mol Pharmacol. 2018;94(3):1092–1100. doi:10.1124/MOL.118.112557,

46. Norouzi S, Adulcikas J, Henstridge DC, Sonda S, Sohal SS, Myers S. The zinc transporter zip7 is downregulated in skeletal muscle of insulin-resistant cells and in mice fed a high-fat diet. Cells. 2019;8(7). doi:10.3390/CELLS8070663,

47. Troche C, Aydemir TBTB, Cousins RJRJ, et al. Zinc transporter Slc39a14 regulates inflammatory signaling associated with hypertrophic adiposity. Am J Physiol Endocrinol Metab. 2016;310(4):E258–E268. doi:10.1152/ajpendo.00421.2015

48. Aydemir TB, Troche C, Kim MH, Cousins RJ. Hepatic ZIP14-mediated Zinc Transport Contributes to Endosomal Insulin Receptor Trafficking and Glucose Metabolism. J Biol Chem. 2016;291(46):23939–23951. doi:10.1074/JBC.M116.748632

49. Kim MHMH, Aydemir TBTB, Kim J, Cousins RJRJ. Hepatic ZIP14-mediated zinc transport is required for adaptation to endoplasmic reticulum stress. Proceedings of the National Academy of Sciences. 2017;114(29):E5805–E5814. doi:10.1073/pnas.1704012114

50. Aydemir TBTBTB, Liuzzi JPJP, McClellan S, Cousins RJRJ. Zinc transporter ZIP8 (SLC39A8) and zinc influence IFN-γ expression in activated human T cells. J Leukoc Biol. 2009;86(2):337–348. doi:10.1189/jlb.1208759

51. Liuzzi JP, Lichten LA, Rivera S, et al. Interleukin-6 regulates the zinc transporter Zip14 in liver and contributes to the hypozincemia of the acute-phase response. Proc Natl Acad Sci U S A. 2005;102(19):6843–6848. doi:10.1073/pnas.0502257102

52. Mitchell SB, Hung YH, Thorn TL, et al. Sucrose-induced hyperglycemia dysregulates intestinal zinc metabolism and integrity: risk factors for chronic diseases. Front Nutr. 2023;10. doi:10.3389/fnut.2023.1220533

53. Aydemir TB, Sitren HS, Cousins RJ. The zinc transporter Zip14 influences c-met phosphorylation and hepatocyte proliferation during liver regeneration in mice. Gastroenterology. 2012;142(7). doi:10.1053/j.gastro.2012.02.046

54. Aydemir TB, Chang SMM, Guthrie GJ, et al. Zinc Transporter ZIP14 Functions in Hepatic Zinc, Iron and Glucose Homeostasis during the Innate Immune Response (Endotoxemia). Nerurkar P V., ed. PLoS One. 2012;7(10):e48679. doi:10.1371/journal.pone.0048679

55. Hung YH, Kim Y, Mitchell SB, Thorn TL, Aydemir TB. Absence of Slc39a14/Zip14 in mouse pancreatic beta cells results in hyperinsulinemia. Am J Physiol Endocrinol Metab. 2024;326(1). doi:10.1152/AJPENDO.00117.2023

56. Martinez-Guryn K, Leone V, Chang EB. Regional Diversity of the Gastrointestinal Microbiome. Cell Host Microbe. 2019;26(3):314–324. doi:10.1016/j.chom.2019.08.011

57. Jensen BAH, Heyndrickx M, Jonkers D, et al. Small intestine vs. colon ecology and physiology: Why it matters in probiotic administration. Cell Rep Med. 2023;4(9). doi:10.1016/j.xcrm.2023.101190

58. Atreya R, Bojarski C, Kühl AA, Trajanoski Z, Neurath MF, Siegmund B. Ileal and colonic Crohn’s disease: Does location makes a difference in therapy efficacy? Current Research in Pharmacology and Drug Discovery. 2022;3. doi:10.1016/j.crphar.2022.100097

59. Ammendola S, Pasquali P, Pistoia C, et al. High-affinity Zn2+ uptake system ZnuABC is required for bacterial zinc homeostasis in intracellular environments and contributes to the virulence of Salmonella enterica. Infect Immun. 2007;75(12):5867–5876. doi:10.1128/IAI.00559-07,

60. Reed S, Neuman H, Moscovich S, Glahn RP, Koren O, Tako E. Chronic zinc deficiency alters chick gut microbiota composition and function. Nutrients. 2015;7(12):9768–9784. doi:10.3390/nu7125497

61. Chen X, Jiang Y, Wang Z, et al. Alteration in Gut Microbiota Associated with Zinc Deficiency in School-Age Children. Nutrients. 2022;14(14). doi:10.3390/NU14142895

62. Huang Z, Wang C, Huang Q, Yan Z, Yin Z. Hungatella hathewayi impairs the sensitivity of colorectal cancer cells to 5-FU through decreasing CDX2 expression. Hum Cell. 2023;36(6):2055–2065. doi:10.1007/S13577-023-00938-Y/FIGURES/6

63. Santiago A, Hann A, Constante M, et al. Crohn’s disease proteolytic microbiota enhances inflammation through PAR2 pathway in gnotobiotic mice. Gut Microbes. 2023;15(1):2205425. doi:10.1080/19490976.2023.2205425

64. Pavlidis P, Tsakmaki A, Treveil A, et al. Cytokine responsive networks in human colonic epithelial organoids unveil a molecular classification of inflammatory bowel disease. Cell Rep. 2022;40(13):111439. doi:10.1016/J.CELREP.2022.111439

65. Collins SM. Interrogating the Gut-Brain Axis in the Context of Inflammatory Bowel Disease: A Translational Approach. Inflamm Bowel Dis. 2020;26(4):493–501. doi:10.1093/IBD/IZAA004

66. Bourgonje AR, Meringer H, Roudko V, et al. P0126 Single-cell analysis reveals enrichment of mucosal Th17-polarized T cells in Acute Severe Ulcerative Colitis. J Crohns Colitis. 2025;19(Supplement_1):i511–i512. doi:10.1093/ECCO-JCC/JJAE190.0300

67. Liu L, Xu M, Lan R, et al. Bacteroides vulgatus attenuates experimental mice colitis through modulating gut microbiota and immune responses. Front Immunol. 2022;13:1036196. doi:10.3389/FIMMU.2022.1036196/BIBTEX

68. Kim JS, Kim HK, Lee J, et al. Inhibition of CD82 improves colitis by increasing NLRP3 deubiquitination by BRCC3. Cell Mol Immunol. 2023;20(2):189–200. doi:10.1038/S41423-022-00971-1;SUBJMETA=1726,250,2504,304,342,631,80;KWRD=MECHANISMS+OF+DISEASE,PHAGOCYTES

69. Gilliland A, Chan JJ, De Wolfe TJ, Yang H, Vallance BA. Pathobionts in Inflammatory Bowel Disease: Origins, Underlying Mechanisms, and Implications for Clinical Care. Gastroenterology. 2024;166(1):44–58. doi:10.1053/J.GASTRO.2023.09.019

70. Yan D, Ye S, He Y, et al. Fatty acids and lipid mediators in inflammatory bowel disease: from mechanism to treatment. Front Immunol. 2023;14:1286667. doi:10.3389/FIMMU.2023.1286667/XML/NLM

71. Agouridis AP, Elisaf M, Milionis HJ. An overview of lipid abnormalities in patients with inflammatory bowel disease. Annals of Gastroenterology : Quarterly Publication of the Hellenic Society of Gastroenterology. 2011;24(3):181. Accessed July 1, 2025. https://pmc.ncbi.nlm.nih.gov/articles/PMC3959314/

72. Uehara M, Inoue T, Hase S, Sasaki E, Toyoda A, Sakakibara Y. Decoding host-microbiome interactions through co-expression network analysis within the non-human primate intestine. mSystems. 2024;9(5). doi:10.1128/MSYSTEMS.01405-23/SUPPL_FILE/MSYSTEMS.01405-23-S0001.DOCX

73. Li Z, Xiong W, Liang Z, et al. Critical role of the gut microbiota in immune responses and cancer immunotherapy. Journal of Hematology & Oncology 2024 17:1. 2024;17(1):1–31. doi:10.1186/S13045-024-01541-W

74. Wasana WP, Senevirathne A, Nikapitiya C, et al. A novel pseudoalteromonas xiamenensis marine isolate as a potential probiotic: Anti-inflammatory and innate immune modulatory effects against thermal and pathogenic stresses. Mar Drugs. 2021;19(12):707. doi:10.3390/MD19120707/S1

75. Alam K, Mazumder A, Sikdar S, et al. Streptomyces: The biofactory of secondary metabolites. Front Microbiol. 2022;13:968053. doi:10.3389/FMICB.2022.968053/XML/NLM

76. Cui Y, Zhang L, Wang X, et al. Roles of intestinal Parabacteroides in human health and diseases. FEMS Microbiol Lett. 2022;369(1):1–11. doi:10.1093/FEMSLE/FNAC072

77. Bolourian A, Mojtahedi Z. Streptomyces, shared microbiome member of soil and gut, as “old friends” against colon cancer. FEMS Microbiol Ecol. 2018;94(8). doi:10.1093/FEMSEC/FIY120,

78. Roda G, Chien Ng S, Kotze PG, et al. Crohn’s disease. Nature Reviews Disease Primers 2020 6:1. 2020;6(1):1–19. doi:10.1038/s41572-020-0156-2

79. Gao HL, Zheng W, Xin N, et al. Zinc deficiency reduces neurogenesis accompanied by neuronal apoptosis through caspase-dependent and -independent signaling pathways. Neurotox Res. 2009;16(4):416–425. doi:10.1007/S12640-009-9072-7,

80. Eide DJ. The oxidative stress of zinc deficiency. Metallomics. 2011;3(11):1124–1129. doi:10.1039/C1MT00064K

81. Mitchell SB, Thorn TL, Lee MT, et al. Metal transporter SLC39A14/ZIP14 modulates regulation between the gut microbiome and host metabolism. Am J Physiol Gastrointest Liver Physiol. Published online October 24, 2023. doi:10.1152/AJPGI.00091.2023

82. Das NK, Schwartz AJ, Barthel G, et al. Microbial Metabolite Signaling Is Required for Systemic Iron Homeostasis. Cell Metab. 2020;31(1):115–130.e6. doi:10.1016/j.cmet.2019.10.005

83. Johnson EL, Heaver SL, Waters JL, et al. Sphingolipids produced by gut bacteria enter host metabolic pathways impacting ceramide levels. Nat Commun. 2020;11(1). doi:10.1038/s41467-020-16274-w

84. Ewels P, Magnusson M, Lundin S, Käller M. MultiQC: summarize analysis results for multiple tools and samples in a single report. Bioinformatics. 2016;32(19):3047–3048. doi:10.1093/BIOINFORMATICS/BTW354

