## Supplemental Figures for "Multiomics Approach to Regionally Profile Zinc-Driven Host-Gut Microbiome Interactions in the Intestinal Tract"

**Supplementary Figures**

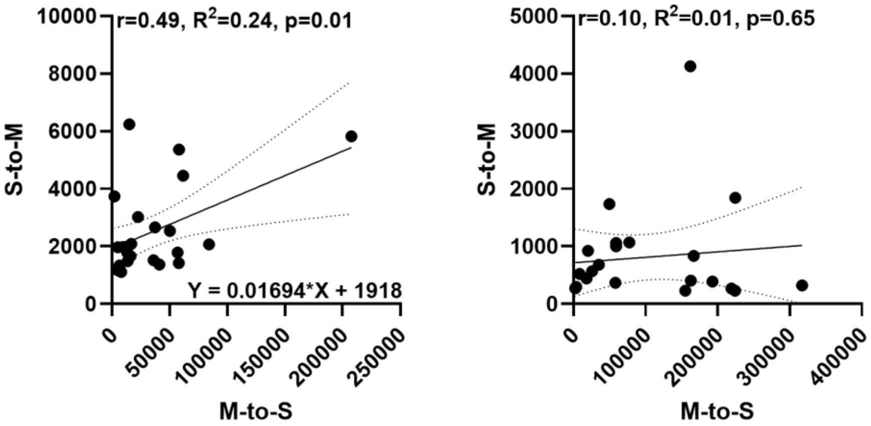


**Supplementary Figure 1.** Correlations between the luminal zinc contents. $^{65}Zn$ radioisotope tracing was conducted in both the serosal-to-mucosal (Subcutaneous injection) and mucosal-to-mucosal (Oral gavage) directions. Radioactivity was measured by gamma counting 3 hours after $^{65}Zn$ administration in the lumen of the small intestine and colon. Simple linear regression plots were generated for the small intestine (A) and the colon (B). M-to-S: Mucosal-to-serosal; S-to-M: Serosal-to-mucosal.

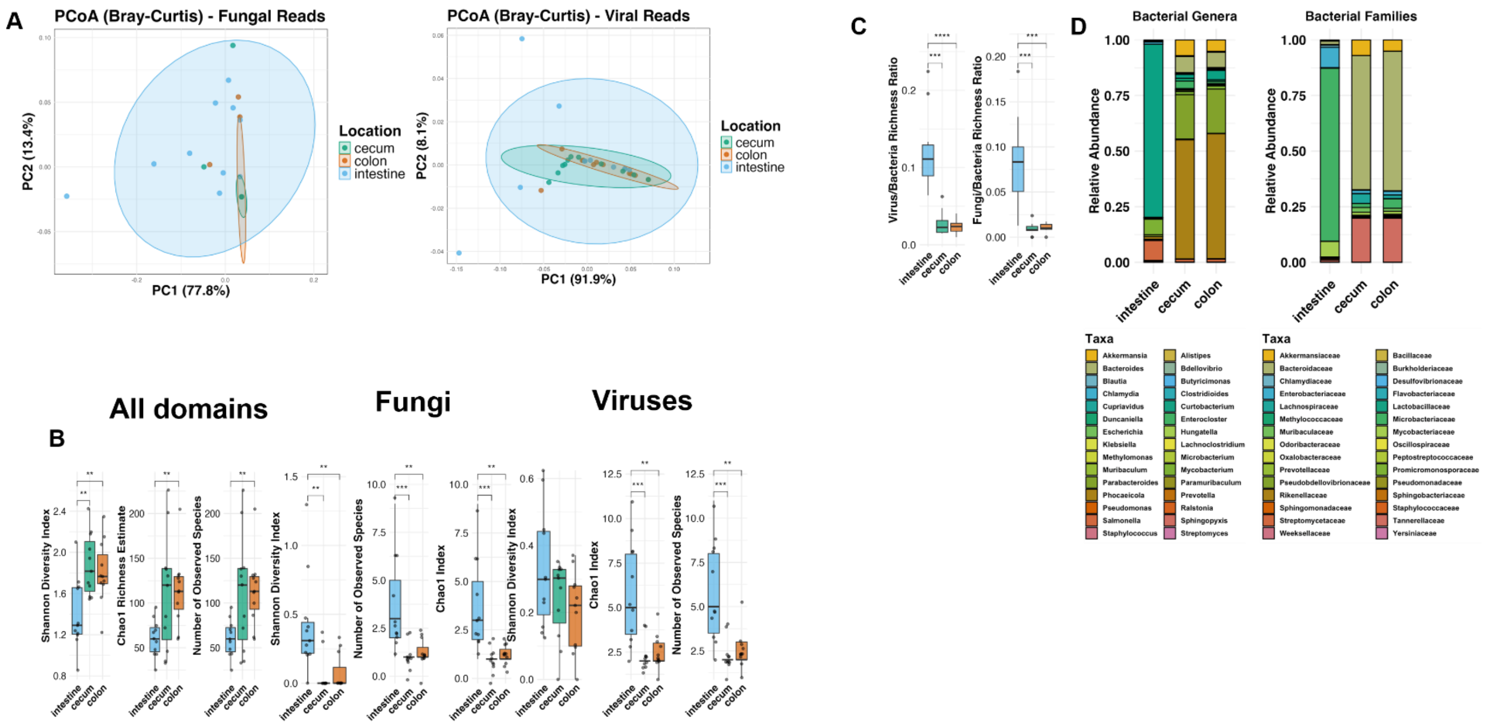


**Supplementary Figure 2.** (A) PcoA plot of fungal and viral reads by intestinal region. Principal Coordinates Analysis (PCoA) based on Bray-Curtis dissimilarity was performed on fungal (left) and viral (right) metagenomic reads derived from the Kraken2 PlusPF database. Analyses were conducted in R version 4.4.2 using the vegan, ggplot2, and phyloseq packages. Each point represents a sample, colored by intestinal region (cecum, colon, intestine), and ellipses indicate 95% confidence intervals for group clustering. (B) Alpha diversity of microbial communities across intestinal regions for all domains, fungi, and viruses. Boxplots represent three alpha diversity metrics: Shannon diversity index, Chao1 richness estimate, and number of observed species calculated from metagenomic reads and separated by intestinal region (intestine, cecum, colon). The panel shows diversity across all domains (bacteria, fungi, viruses) and diversity for fungal and viral reads specifically, respectively. Statistically significant pairwise comparisons were assessed using the Wilcoxon rank-sum test and are indicated as follows: $p < 0.05$ (*), $p < 0.01$ (**), $p < 0.001$ (***). (C) Relative richness of fungal and viral communities compared to bacteria across intestinal regions. Boxplots show the ratios of fungal-to-bacterial richness (right) and viral-to-bacterial richness (left) across intestinal regions (intestine, cecum, colon). Richness values were calculated for each sample based on the number of observed species identified using Bracken2-corrected outputs from Kraken2. Ratios were computed by dividing fungal or viral richness by bacterial richness for each corresponding sample. Pairwise statistical comparisons were performed using the Wilcoxon rank-sum test. **$p < 0.001$, *$p < 0.0001$. (D) Relative abundance of bacterial families and genera across intestinal regions. Stacked bar charts showing the mean relative abundance of the top 30 bacterial genera (left) and families (right) across the small intestine, cecum, and colon. Taxa were ranked by cumulative abundance across all samples, and only the 30 most abundant taxa across the full dataset were retained for plotting. Abundances were normalized within each sample and then averaged across all samples from each intestinal location.

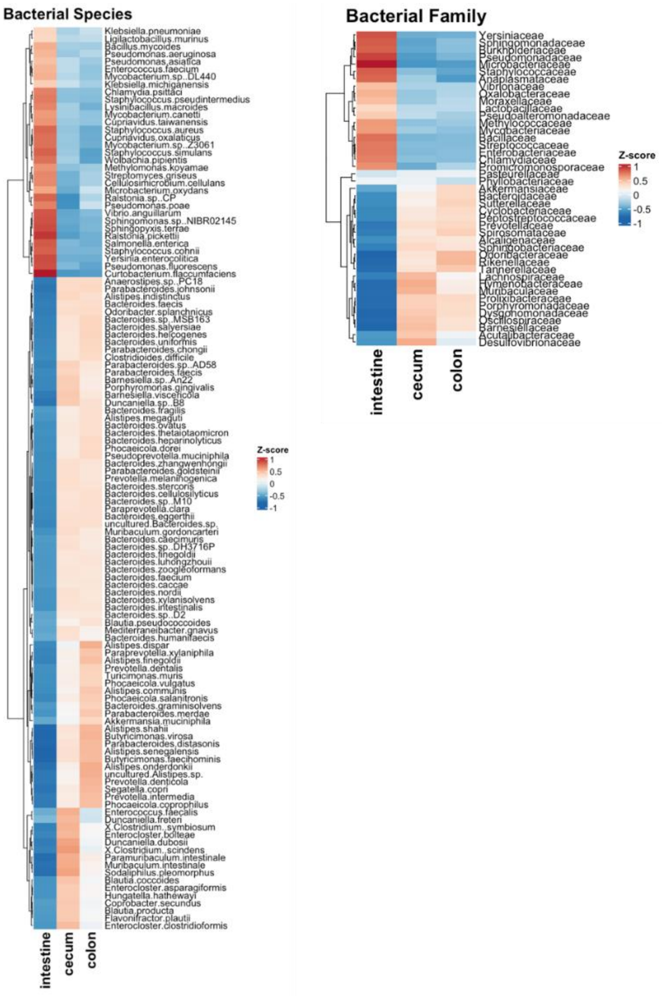


**Supplementary Figure 3. Heatmaps of Differentially Abundant Bacterial Taxa Across Intestinal Regions.** Heatmaps showing Z-score normalized relative abundance of differentially abundant bacterial families (right) and species (left) across the intestine, cecum, and colon in averaged ZnD and ZnA samples. Taxa were identified as significantly differentially abundant (Maaslin2, $p < 0.05$) and Z-scores were computed per taxon across all samples and then averaged within each location to highlight spatial patterns. Clustering of taxa was performed using hierarchical clustering based on Z-scores, while columns were ordered by gut region. Higher Z-scores reflect increased abundance relative to that taxon's overall mean across all samples.

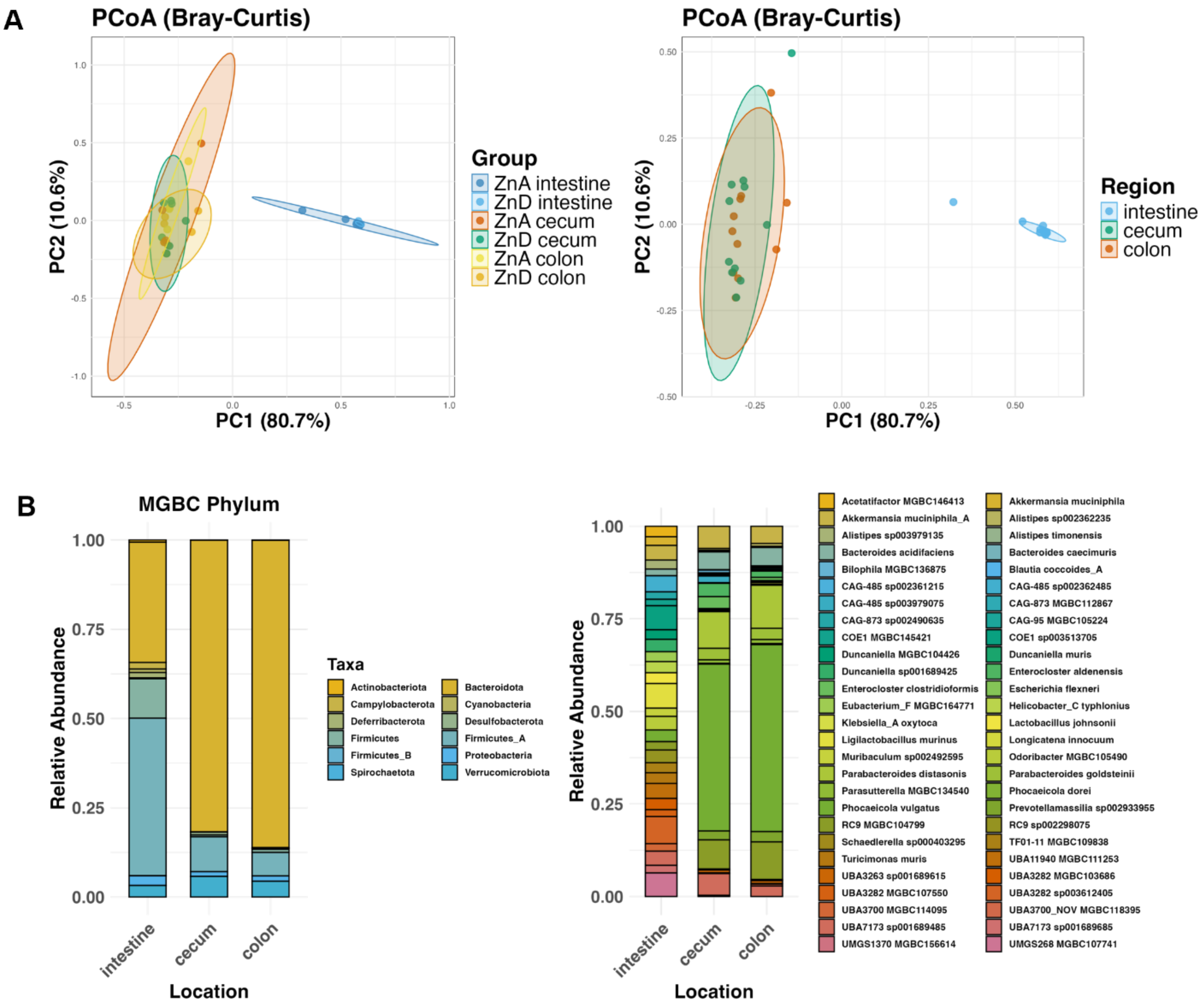


**Supplementary Figure 4. Spatial Microbial Profiling Using the Mouse Gastrointestinal Bacteria Catalogue (MGBC).** (A) PCoA and taxonomic barplots comparing microbial community structure across intestinal regions using the mouse-specific MGBC database. PCoA plots based on Bray-Curtis dissimilarity of MGBC-derived bacterial species-level profiles. Left: samples colored by treatment and region; right: samples colored by region only. (B) Stacked barplots of relative abundance for each gut region at the phylum level (left) and species level (right), averaged across samples. Raw MGBC counts were normalized to relative abundances. Mean relative abundances were then calculated across all samples within each location.

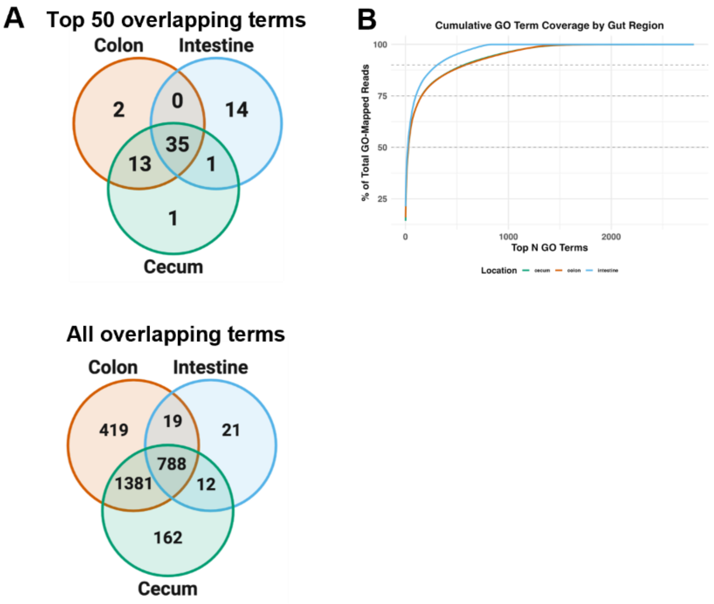


**Supplementary Figure 5.** (A) Overlap of functional annotations across gut regions based on Gene Ontology (GO) terms. Venn diagrams display the number of overlapping GO terms across the colon, intestine, and cecum. The left panel represents all GO terms detected from HUMAnN3 pathway annotation, while the right panel shows the top 50 most abundant GO terms per region based on cumulative read counts. GO term rankings were based on TSS-normalized read counts, and overlaps were visualized using the VennDiagram package in R. (B) Cumulative Contribution of GO Terms to Functional Read Coverage Across Gut Regions. Cumulative coverage curves of GO terms ranked by abundance across the colon, intestine, and cecum. GO terms were derived from HUMAnN3 pathway outputs and filtered to include terms with ≥10 counts per million (CPM) in at least one region and presence in at least two regions. For each location, GO terms were ranked by total mapped reads, and the cumulative percentage of reads was plotted. Dashed lines indicate 50%, 75%, and 90% coverage thresholds.

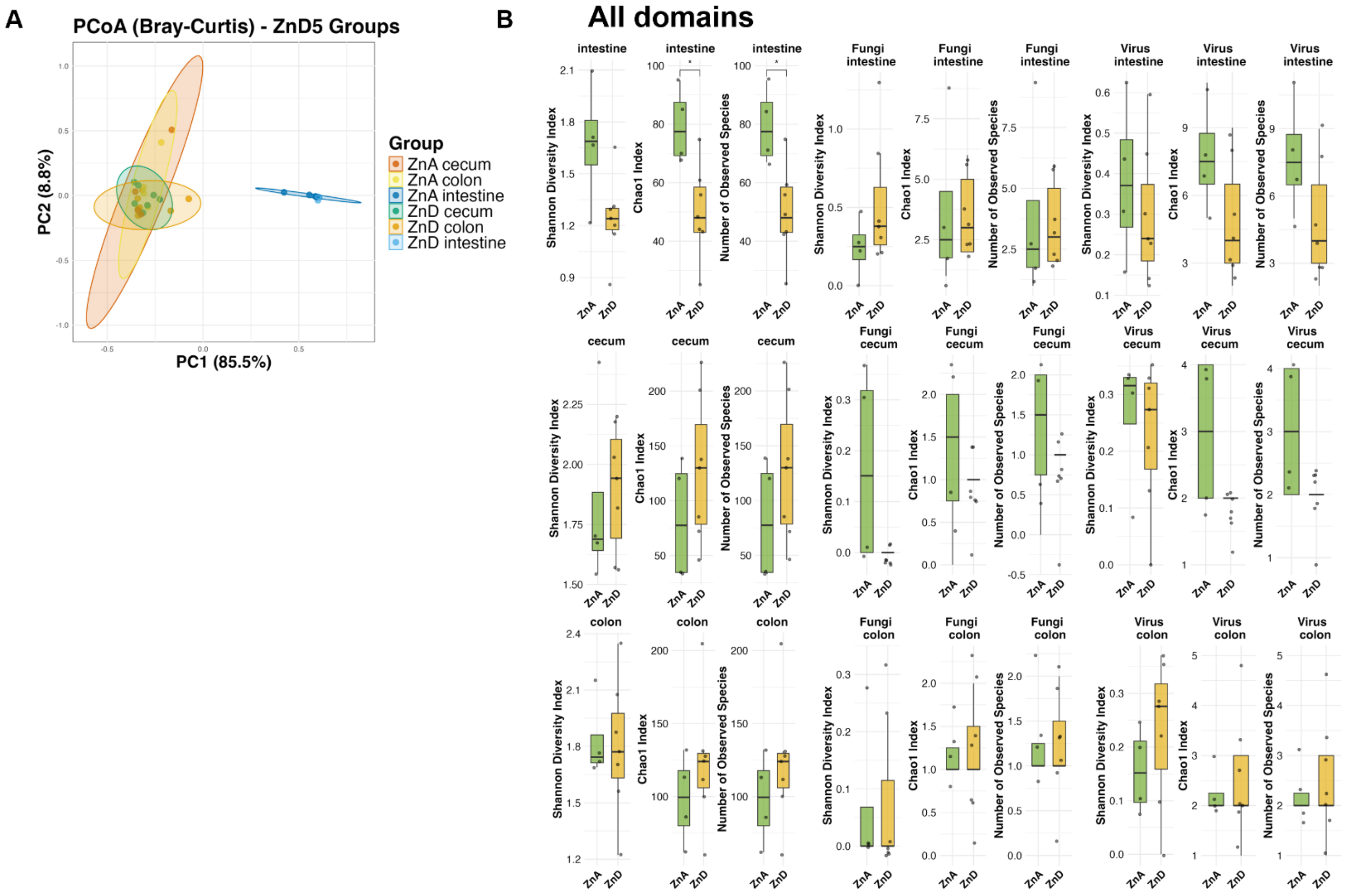


**Supplementary Figure 6.** (A) Spatial Microbial Profiling Using the PlusPF Database. PCoA and taxonomic barplots comparing microbial community structure across intestinal regions using the PlusPF database. PCoA plots based on Bray-Curtis dissimilarity of PlusPF-derived bacterial species-level profiles show strong region-specific clustering of samples. Samples are colored by treatment and region. (B) Alpha Diversity Across Gut Regions and Microbial Kingdoms. Boxplots show alpha diversity metrics (Shannon Diversity Index, Chao1 Index, and observed species) for all domains, fungal, and viral communities across the small intestine, cecum, and colon under ZnA and ZnD) conditions. Diversity was assessed using phyloseq on TSS-normalized Bracken read counts classified with the PlusPF database. Significant differences between treatments were calculated using Wilcoxon rank-sum tests and visualized with p-values ($p < 0.05$).

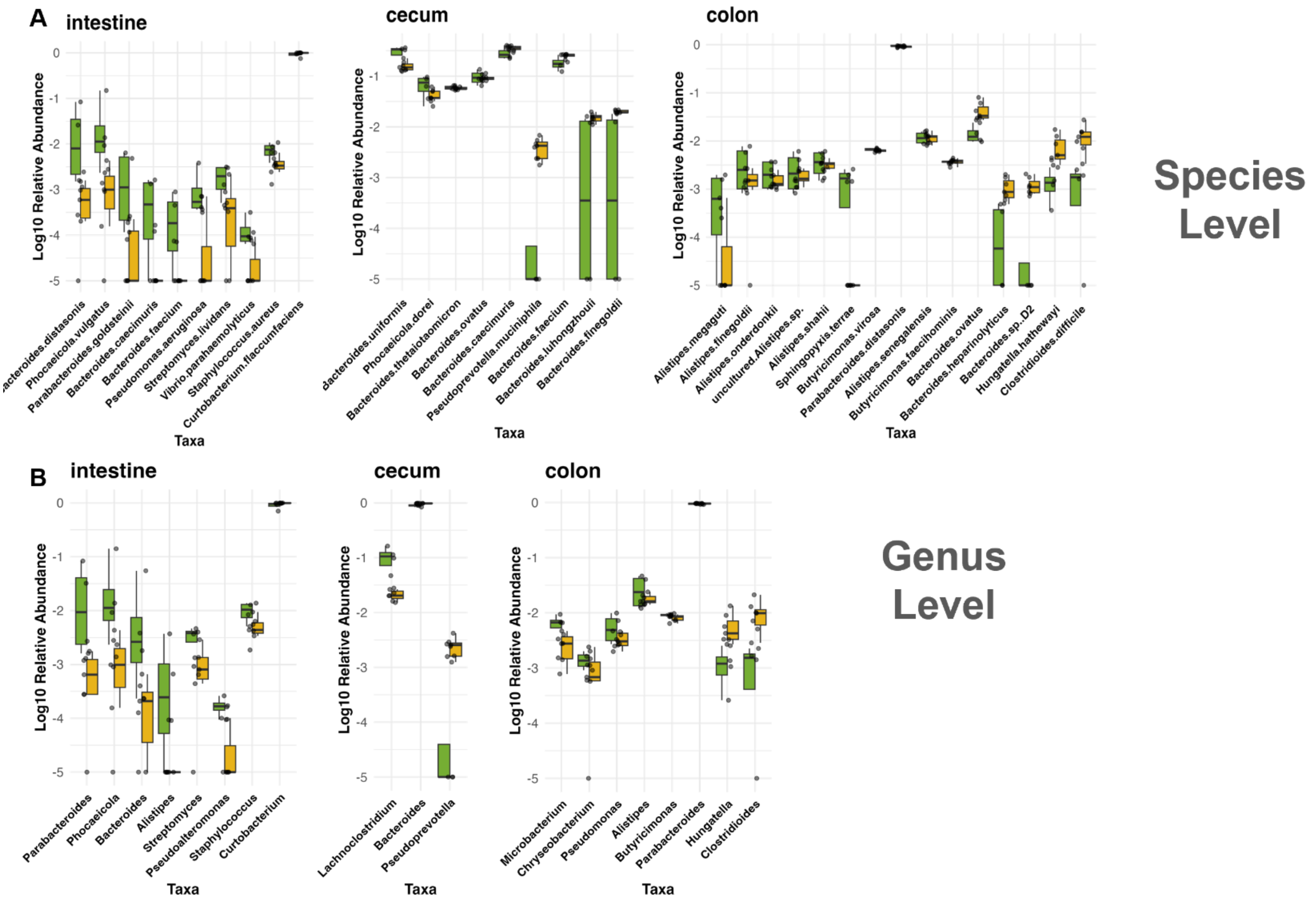


**Supplementary Figure 7.** Boxplots depict differentially abundant bacterial taxa identified between ZnA (green) and ZnD (yellow) conditions across three intestinal regions: the intestine, cecum, and colon at the species and genus levels. Taxa shown were filtered based on significance ($p < 0.05$) from MaAsLin2 differential expression analysis and are plotted using total sum scaling (TSS)-normalized relative abundances ($\log_{10}$-transformed). Only taxa with nonzero expression in both treatment groups were included. Taxa are ordered by $\log_2$ fold change, with the most ZnD-enriched taxa on the right and ZnA-enriched on the left within each plot.

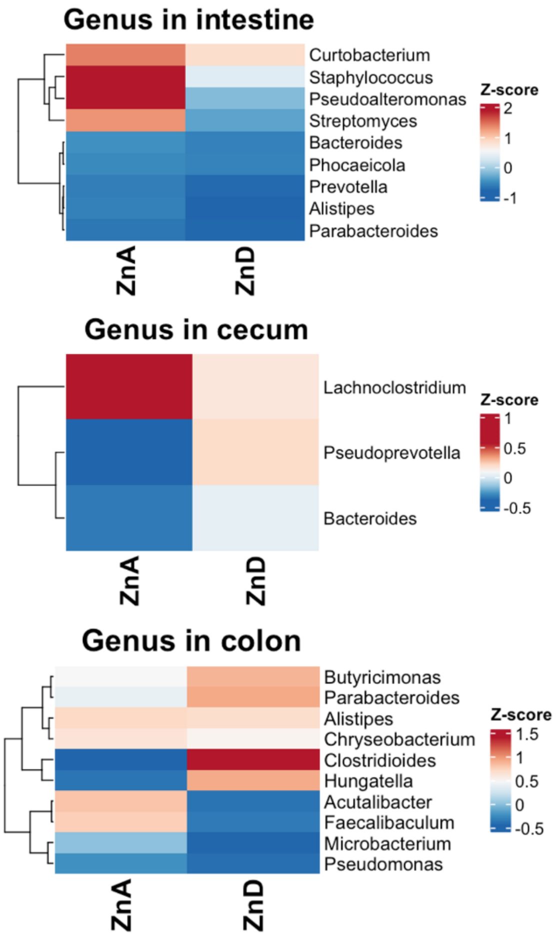


**Supplementary Figure 8. Heatmaps of Differentially Abundant Bacterial Taxa Across Intestinal Regions and Treatment Groups.** Heatmaps showing Z-score normalized relative abundance of differentially abundant bacterial genera in the intestine, cecum, and colon. Taxa were identified as significantly differentially abundant (Maaslin2, $p < 0.05$) using ZnA as the reference for the treatment group. Z-scores were computed across all samples and then averaged by treatment group within each region to highlight spatial patterns. Clustering of taxa was performed using hierarchical clustering based on Z-scores, while columns were ordered by treatment. A higher Z-score indicates higher relative abundance compared to the taxon's overall mean across the two treatments.

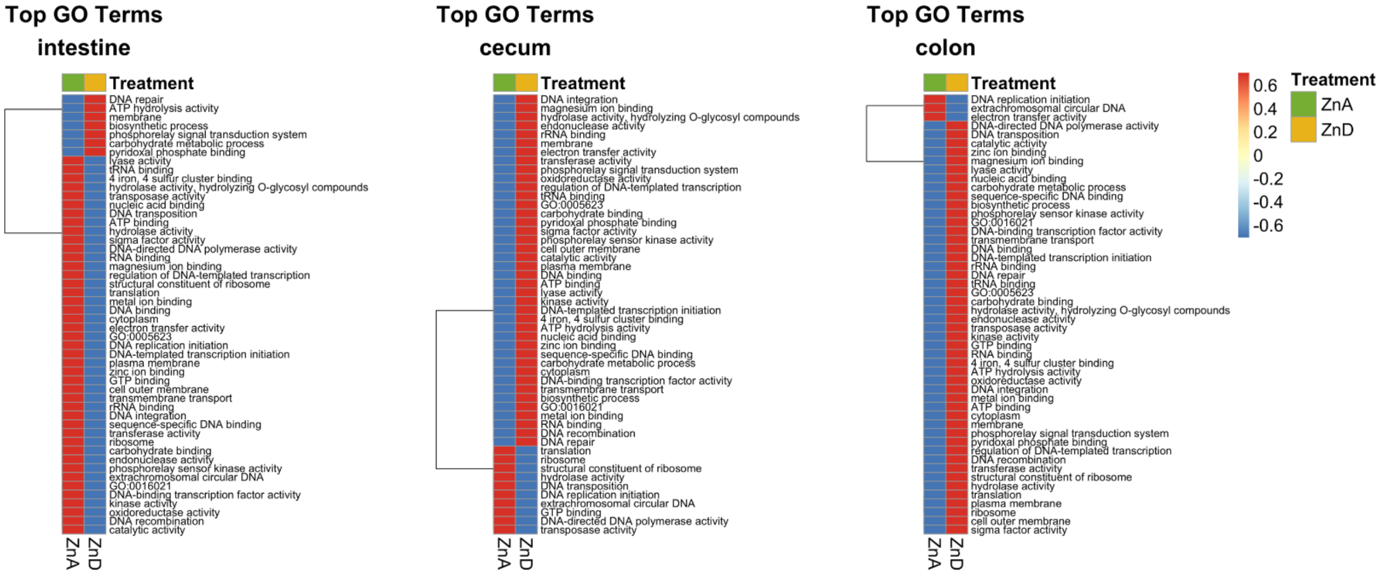


**Supplementary Figure 9. Zinc-dependent spatial remodeling of microbial functional pathways across the intestine, cecum, and colon.** Heatmaps display the top 50 most variable GO terms by location based on HUMAnN3-annotated metagenomic profiles from ZnA and ZnD conditions. For each anatomical region, mean relative abundances were computed per treatment group after filtering out GO terms not detected in both ZnA and ZnD samples. Rows represent GO terms and columns represent treatment groups (ZnA, ZnD), with values Z-score normalized across each row, highlighting directional changes. Hierarchical clustering (Euclidean distance, complete linkage) was applied to GO terms.

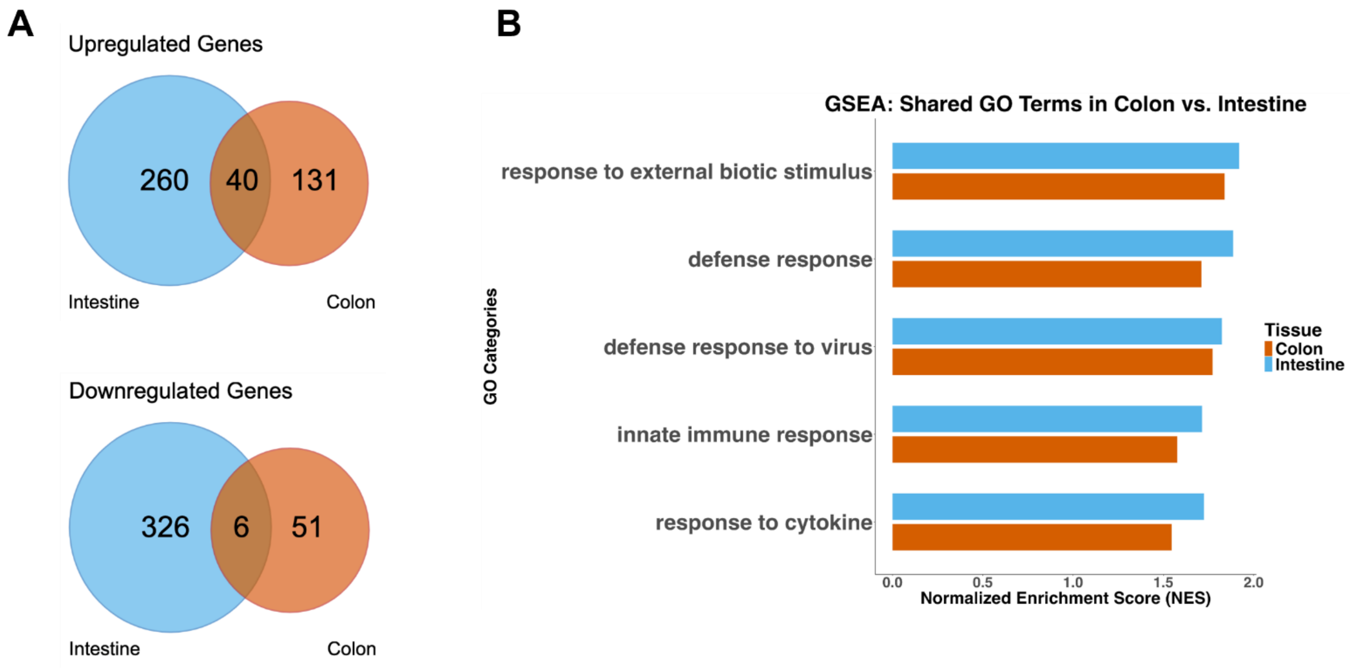


**Supplementary Figure 10. Shared GO terms between the intestine and colon include immune function pathways.** (A) Common and unique upregulated and downregulated genes between the small intestine (blue) and colon (red) tissues in ZnD-fed mice. (B) Enrichment scores for the intestine and colon within the top 5 shared GO terms ($p < 0.05$).
